# Rhodo-Box: a Synthetic Biology Toolbox to Facilitate Metabolic Engineering of *Rhodobacter sphaeroides*

**DOI:** 10.1101/2025.10.31.685836

**Authors:** Matic Kostanjšek, Antoine Raynal, George Dimopoulos, Gerrich Behrendt, Vitor A.P. Martins dos Santos, Jules Beekwilder, Christos Batianis, Ruud A. Weusthuis, Enrique Asin-Garcia, Markus M.M. Bisschops

## Abstract

*Rhodobacter sphaeroides* is a purple non-sulphur alphaproteobacterium with a highly versatile metabolism. This microorganism holds promise as a chassis for sustainable biomanufacturing of numerous chemicals. Yet, its potential is constrained by a lack of standardized, well-characterized genetic elements to tune gene expression such as transcriptional promoters and ribosome binding sites (RBSs). In this study, we present Rhodo-Box, a comprehensive toolkit for *R. sphaeroides* created by adapting and extending the Zymo-Parts modular cloning framework. Using Rhodo-Box we built and characterized: (a) three broad-host origins of replication (pBBR1, RK2 and RSF1010), (b) a set of 13 promoters, (c) four inducible expression systems (NahR-P*_salTTC_*, LacI-P*_lacT7A1_O3O4_*, VanR-P*_vanCC_*, and XylS-P_m_), (d) 11 RBSs, and (e) four transcriptional terminators. Furthermore, we present a semi-automated, user-friendly cloning approach which enables rapid construction of *R. sphaeroides* strains. The Rhodo-Box toolkit equips *R. sphaeroides* with a standardized, automation-compatible collection of parts and workflows essential for efficient design–build–test–learn cycles and advanced metabolic engineering.

**Graphical abstract:** 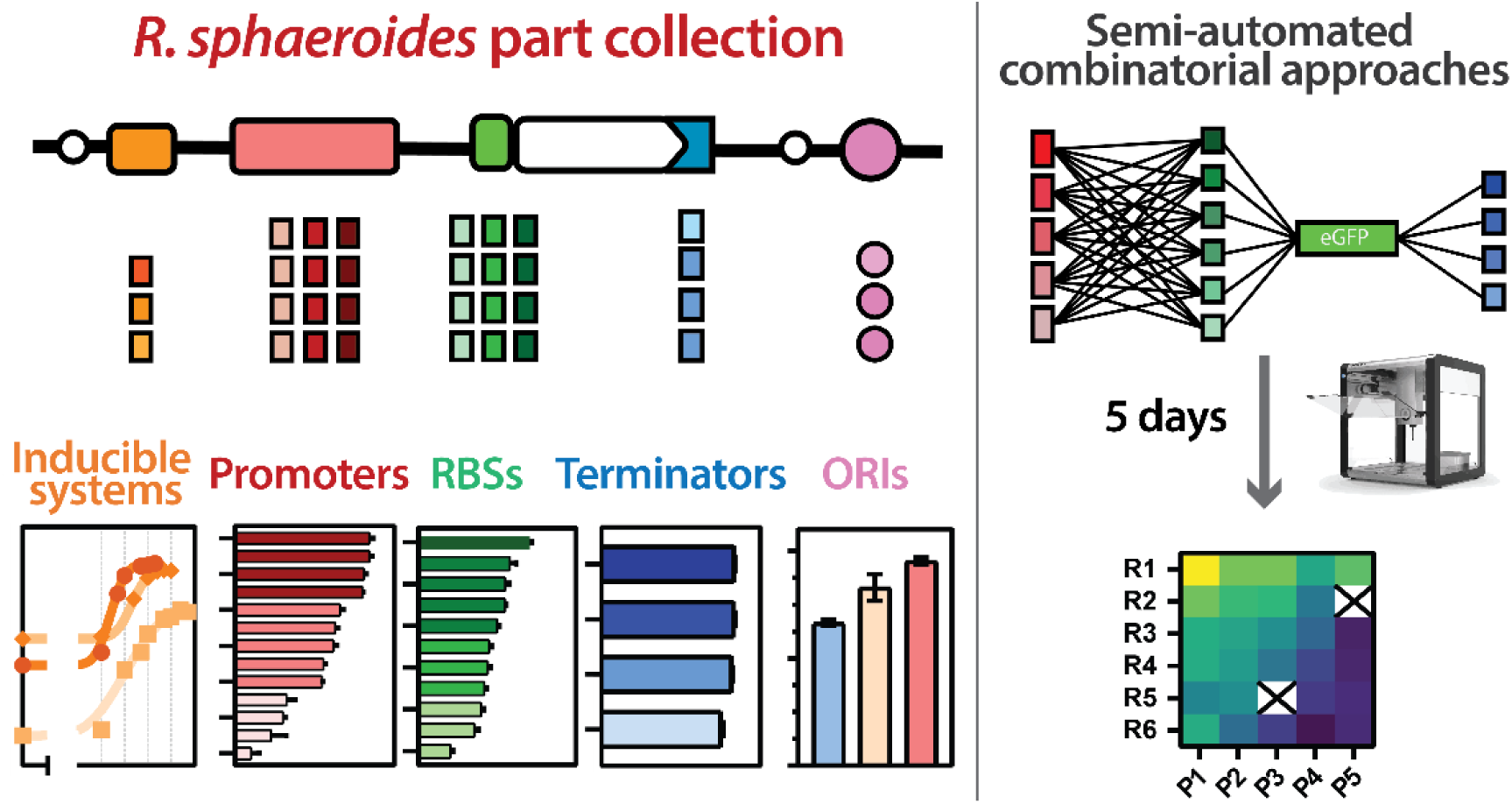

## Introduction

*Rhodobacter sphaeroides* has emerged as a promising cell factory for the sustainable biomanufacturing of numerous chemicals. Its significant metabolic versatility favors extensive rewiring of endogenous pathways, dramatically enhancing its natural ability to produce high-value compounds (Beekwilder et al., 2014; Chen et al., 2024; Hu et al., 2021; Lee et al., 2020; Lu et al., 2015; Orsi et al., 2020). *R. sphaeroides’* industrial potential, characterized by its production metrics (titers, rates, yields), could be further improved with a deeper embedding in the synthetic biology DBTL (design-build-test-learn) framework (Orsi et al., 2021).

The implementation of a comprehensive and standardized synthetic biology toolbox in *R. sphaeroides,* that enables predictable and scalable strain engineering, would improve and diversify its scope of biotechnological applications. Although a BioBrick-compatible vector system was developed for this organism (Lu et al., 2014; Tikh et al., 2014), modular cloning-based assembly systems (MoClo) have since emerged as more effective alternatives in various industrially relevant microorganisms (Behrendt et al., 2022; Damalas et al., 2020; Guiziou et al., 2016; Lee et al., 2015). To date, no specific MoClo-based toolbox has been reported for *R. sphaeroides,* representing a major bottleneck for systematic metabolic engineering in this promising organism.

Among existing platforms, the Zymo-Parts modular cloning toolbox developed for *Zymomonas mobilis* by Behrendt et al. (2022) is of particular interest (Figure 1). So far, this toolbox has been mostly utilized for applications in *Z. mobilis* and *E. coli* (Frohwitter et al., 2024; Wichmann et al., 2023). It features a double loop (“Golden Braid”-like) architecture which enables the recursive and infinite assembly of transcriptional units within levels 1 and 2. Furthermore, the toolbox introduces the modular and hierarchical construction of polycistronic operons, allowing for simultaneous control over the expression of several genes within the same transcriptional unit. Due to these advantages, the Zymo-Parts framework was selected as the foundation for developing a synthetic biology toolbox for *R. sphaeroides*.

**Figure 1:**
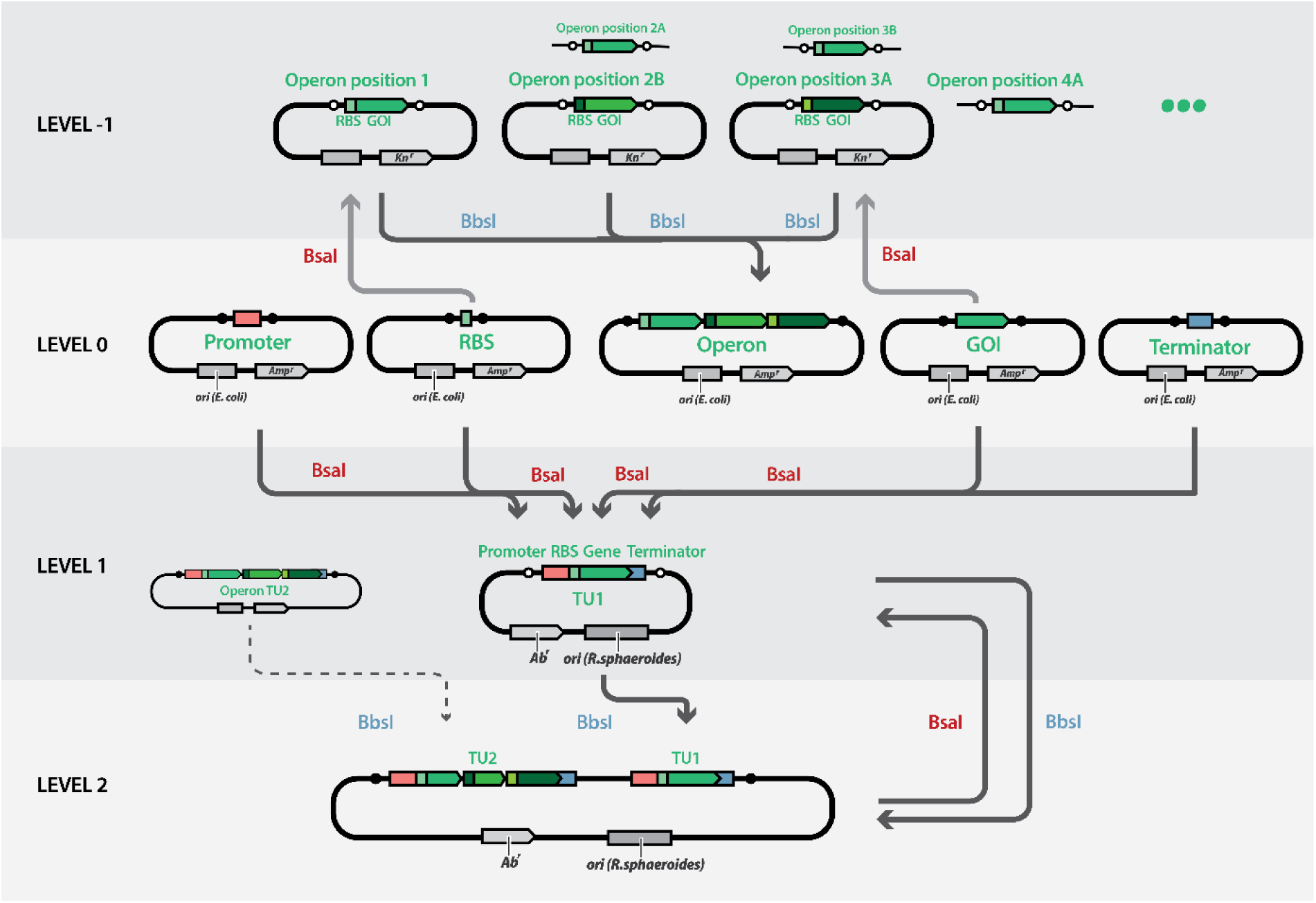
Graphical representation of the Zymo-Parts cloning system developed by Behrendt et al. (2022). The system is composed of four main levels. Level 0 is the collection of individual, basic parts. A gene of interest (GOI) and RBS can be assembled into level -1 to create operon fragments, which can be assembled back into polycistronic operon parts in level 0. Level 0 parts can be assembled to transcriptional units in level 1. By constructing specific positioned level 1 plasmids, multiple transcriptional units can be assembled together into a level 2 plasmid. Double loop architecture then allows for recursive extension of the plasmids within iterative level 1 and level 2 assemblies.

Synthetic biology toolboxes strongly rely on collections of well-characterized genetic parts, such as RBSs, transcriptional promoters (including inducible systems), transcription terminators and origins of replications (ORIs). Such quantitatively defined part libraries are essential for implementing sophisticated approaches in metabolic engineering (Moreno-Paz et al., 2024; Radivojević et al., 2020).

To date, the characterization of only a limited set of genetic parts, including promoters, RBSs and terminators, has been reported for *R. sphaeroides*. Huo (2011) designed a comprehensive BioBrick-compatible expression system. However, only native promoters were assessed which are likely subject to native regulation (Henry et al., 2020), and the synthetic RBSs were constructed by varying a single reference sequence, which can introduce design bias and limit the dynamic range of the RBSs (Höllerer et al., 2020; Huo, 2011). More recently, Shi et al. (2021) developed a promoter library based on mutated variants of a native promoter. However, the characterized promoter parts retained the native RBS, which compromises their modularity. Detailed characterization of orthogonal elements, synthetic or heterologous, is lacking and could greatly expand the dynamic ranges for gene expression in *R. sphaeroides*.

Inducible expression systems offer another critical way to control gene expression. In *R. sphaeroides*, promoters induced by crystal violet (CV), isopropyl β-D-1-thiogalactopyranoside (IPTG), oxygen or light have been utilized (Hall et al., 2024; Ind et al., 2009; Kretz et al., 2023; Tikh et al., 2014). However, each of these systems has significant limitations: CV is cytotoxic and can negatively impact cell growth (Kretz et al., 2023), while oxygen and light are condition-specific inducers, which makes them less suitable when a general, flexible, condition-agnostic and metabolism-neutral inducer is required. Furthermore, the well-established IPTG-based *Lac* promoter functions poorly in *R. sphaeroides* and required extensive engineering efforts to achieve adequate performance (Hall et al., 2024; Ind et al., 2009; Kretz et al., 2023). Therefore, expanding the inducible expression system collection further would enable novel combinatorial approaches that require simultaneous use of several well-characterized and titratable systems (Lee et al., 2024; Meyer et al., 2019).

The use of standardized parts in a molecular toolbox supports modularity, a critical feature for transitioning to automated workflows (Chao et al., 2017; Weber et al., 2016). With a larger collection of standardized elements, the potential design space grows exponentially (Smanski et al., 2014). Manually assembling and screening this space is both time- and labor-intensive. Automation can alleviate these burdens by enabling the construction of a larger portion of the design space while reducing the required hands-on time (Carbonell et al., 2018; Stephenson et al., 2023). Compared to random libraries, design-specific libraries yield higher information content upon screening. In today’s convergence of synthetic biology and ML, such data are valuable for training and refining ML-driven metabolic models (Zampieri et al., 2019). To democratize access to automation, especially for laboratories that lack the financial or technical capacity for large-scale robotic platforms, synthetic biology toolboxes should ideally be compatible with low-cost, user-friendly automated platforms.

In this study, we present Rhodo-Box – a comprehensive synthetic biology toolbox with an extensive collection of characterized genetic elements tailored for *R. sphaeroides.* We systematically assessed multiple sets of genetic elements, including 13 constitutive promoters, 4 inducible systems, 11 RBSs, 4 transcriptional terminators, and 3 origins of replication (ORIs). We show broad dynamic ranges– 250-fold for promoters and 49-fold for RBSs. Furthermore, the inducible systems exhibited a low basal expression and high tunability, providing versatility for strain engineering designs. Finally, we also established an accessible semi-automated cloning workflow to build *R. sphaeroides* libraries in a time-efficient manner using an Opentrons OT2 platform.

## Materials and Methods

### Strains and media

*R. sphaeroides* 265-9c and *E. coli* strains DH5α and ST18 were used in this study (Table S1). All plasmids used in this study can be found in the supporting information section (Tables S2). *E. coli* DH5α was used to propagate the plasmids and was cultivated in LB medium (10 g/L tryptone, 5 g/L yeast extract, 5 g/L NaCl) at 37 °C. When required, 100 µg/mL of ampicillin, 50 µg/mL of kanamycin, 10 µg/mL of gentamicin, 100 µg/mL of spectinomycin, or 25 µg/mL of chloramphenicol was added. *R. sphaeroides* was cultivated in either LB or modified Sistrom’s minimal media (SMM) containing 3.7 g/L (NH_4_)_2_SO_4_, 3.48 g/L KH_2_PO_4_, 0.5 g/L NaCl, 0.3 g/L MgSO_4_·7H_2_O, 0.0334 g/L CaCl_2_·2H_2_O, 0.002 g/L FeSO_4_·7H_2_O, and 0.0002 g/L (NH_4_)_6_Mo_7_O_24_. Trace elements were added (0.1% v/v) from a stock solution containing 1.765 g/L disodium EDTA, 10.95 g/L ZnSO_4_·7H_2_O, 7.2 g/L FeSO_4_·7H_2_O, 1.54 g/L MnSO_4_·7H_2_O, 0.392 g/L CuSO_4_·5H_2_O, 0.248 g/L Co(NO_3_)_2_·6H_2_O, and 0.00114 g/L H_3_BO_3_. Vitamins were added (0.01% v/v) from a stock containing 10 g/L nicotinic acid, 5 g/L thiamine HCl, and 0.107 g/L biotin. Glucose was added as the sole carbon source to a final concentration of either 10 or 3.6 g/L. When cultivating plasmid-carrying strains, 50 µg/mL kanamycin, 10 µg/mL gentamicin, 100 µg/mL spectinomycin, or 25 µg/mL chloramphenicol was added accordingly. All strains were stored at -80 °C in 26 % glycerol.

### Molecular protocols

PCR reactions for domestication of genetic parts were performed using the Q5 high-fidelity polymerase (NEB), according to manufacturer’s instructions. The product was either isolated from agarose gel following a gel electrophoresis or directly from the PCR reaction using the GeneJET Gel Extraction and DNA Cleanup Micro kit (Thermo Scientific). If required, any plasmid template was removed after the PCR reaction by adding 1 µL of DpnI (NEB) enzyme and digesting at 37 °C for 60 min, followed by 10 min of inactivation at 80 °C, prior to product isolation.

The protocols for constructing the level 0 entry vectors with new parts and a comprehensive description of the Zymo-Parts cloning toolbox are found in Supplementary notes 1-4.

To assemble the parts, Golden-Gate assembly was used with either BbsI (NEB) or BsaI-HF (NEB) as the restriction enzyme. Briefly, 1 µL of 10× CutSmart buffer (NEB), 0.5 µL of recombinant bovine serum albumin (rBSA, NEB), 0.5 µL of T4 ligase (NEB), and 0.5 µL of the required restriction enzyme (NEB) were mixed with 1 µL of the selected entry vector, and 1 µL of the toolkit part (plasmid concentrations were diluted to 50-100 ng/µL). The volume was adjusted to 10 µL using Milli-Q H_2_O. For the reaction, 30 cycles of 3 min at 37 °C, followed by 5 min at 16 °C were carried out, and ended with a final digestion of 10 min at 37 °C, and a final inactivation of 10 min at 80 °C.

Diagnostic PCRs were performed using the Phire polymerase 2× Master-Mix (Thermo Scientific), in accordance with manufacturer’s instructions. When performing diagnostic PCRs on *R. sphaeroides* colonies, a small amount of biomass from a colony was first boiled in 20 µL of Milli-Q H_2_O. The suspension was then briefly centrifuged to remove the insoluble cell debris, and 1 µL of the supernatant was used as a template in a 10 µL total volume of the PCR.

### Conjugation and transformation

To propagate assembled plasmids, 5 µL of the Golden Gate reaction was mixed with 50 µL of chemically competent DH5α cells. After 20 min incubation on ice, a heat shock at 42 °C was applied for 30 s before putting on ice for 2 min. 200 µL of LB was then added to the competent cell mixture and the mixture was incubated at 37 °C for 1 h. Finally, 100 µL of the mixture was plated on an LB agar plate with appropriate antibiotics. Plasmids were isolated using the GeneJet plasmid purification MiniPrep kit (Thermo Scientific). 1 µL of isolated level 1 or level 2 plasmids was mixed with 50 µL of chemically competent *E. coli* ST18 cells (Thoma and Schobert, 2009) and transformed using the same protocol described above. 50 µg/mL 5-aminolevolunic acid (5-ALA) was added to the recovery LB and solid LB to complement the auxotrophy of *E. coli* ST18.

Biparental conjugation was performed to transfer plasmids from *E. coli* ST18 strains into *R. sphaeroides* strains. *R. sphaeroides* strains were inoculated from a glycerol stock in 5 mL LB medium and incubated overnight at 30 °C and 250 rpm. The *E. coli* ST18 strain carrying the desired plasmid was streaked on a LB plate supplemented with 50 µg/mL 5-ALA and the appropriate antibiotic and grown overnight at 37 °C. 300 µL of the *R. sphaeroides* culture was mixed with the cells of a single *E. coli* ST18 colony. The mixture was centrifuged at 6,000 x g for 3 min, and cells were concentrated by removing the supernatant and resuspending in 20 µL of fresh LB medium supplemented with 5-ALA. Cells were spotted on an LB plate supplemented with 5-ALA and incubated at 30 °C for 4-5 h. Cells were then resuspended in 300 µL of fresh LB medium and centrifuged to remove all remaining 5-ALA. After removing the supernatant, the pellet was again resuspended in 500 µL of fresh LB media. Finally, 100 µL was plated onto LB with appropriate antibiotic and incubated at 30 °C for 3 d to select for *R. sphaeroides* containing the plasmid.

### Cultivations

A plate reader (BioTek Synergy H1) was used to assess growth and fluorescence intensities of *R. sphaeroides* strains expressing eGFP. To perform the cultivation experiment, the desired strain was first streaked from a glycerol stock onto an LB plate with appropriate antibiotics and incubated at 30 °C for 3 d. A single colony was then transferred into a 50-mL tube containing 5 mL of LB with appropriate antibiotics and cultured overnight at 30°C and 250 rpm. OD_600_ of the precultures was measured and the amount of the inoculum was adjusted so that the second preculture reached an OD_600_ of 1.5 the following day. Strains were inoculated in SMM with appropriate antibiotics and glucose, and cultivated at 30 °C and 250 rpm. OD_600_ of the overnight cultures was measured and an appropriate amount of culture was centrifuged and resuspended in fresh SMM with glucose and appropriate antibiotics to reach an OD_600_ of 0.1. 200 μL of the culture was transferred into each well of a black 96-well plate. Strains were assessed in technical triplicates, and the plate was incubated at 30 °C with double-orbital shaking (amplitude, speed) for 64 h. OD_600_ was measured every 5 min, as well as eGFP fluorescence by measuring the emission at 512 nm after excitation at 488 nm. Fluorescence readings were normalized by dividing by the media-blanked OD_600_ values at the corresponding timepoint. Final biomass-normalized fluorescence values were obtained by finding the highest moving average value of about 2 h during the exponential growth phase of the cultivation. The plots showing the normalized fluorescence during the cultivation and the intervals corresponding to highest average normalized fluorescence are shown in Figures S9-14.

Inducible systems were assessed in technical triplicates using 96-well plates as described above. To assess the VanR-P*_vanCC_*, NahR-P*_salTTC_*, LacI-P*_lacT7A1_O3O4_,* and XylS-P_m_ systems, vanillic acid (0-5000 µM), salicylic acid (0-200 µM), IPTG (0-1000 µM), or 3-methylbenzoate (0-2000 µM) were used as inducers, respectively. The dose-response calculations were performed using GraphPad Prism software (Version 10.3.1, Dotmatics), where the parameters were determined using the Hill equation with variable slope (Y=Bottom + (Top-Bottom)/(1+(EC50/X)^HillSlope). The bottom of the curves was constrained to the mean of normalized fluorescence values of uninduced samples carrying the respective inducible systems.

### Single timepoint measurements

When assessing the constructs with varying ORIs, the *R. sphaeroides* strains were first streaked from a glycerol stock onto an LB plate with appropriate antibiotics and incubated at 30 °C for 3 d. A single colony was then transferred into a 50-mL tube containing 5 mL of LB with appropriate antibiotics and cultured overnight at 30 °C and 250 rpm. OD_600_ of the precultures was measured and the amount of the inoculum was adjusted for the second preculture to reach an OD_600_ of 1.5 the following day. The appropriate volume of the preculture (calculated based on µ = 0.15 h^-1^) was inoculated to 5 mL of SMM (55 mM glucose as sole carbon source) to ensure that the cells were in the mid-exponential growth phase (OD_600_ ≈ 1.5) the following day. Next day, cells were diluted 10x in a Greiner Bio-One black 96-well plate. The OD_600_, and eGFP fluorescence (emission = 512 nm, excitation = 488 nm) were measured in a BioTek Synergy H1 plate reader. Biomass-normalized fluorescence was calculated by dividing the fluorescence values with the media-blanked OD_600_ values.

### Semi-automated workflow

The automated liquid handling machine Opentrons OT2 was used for the development of the semi-automated workflow. Individual automated protocols were built with either the protocol designer web page of Opentrons (https://designer.opentrons.com/), or using the Opentrons Python Protocol API v2.0 (https://docs.opentrons.com/v2/).

Three automated protocols were employed for this workflow. An Excel based Golden Gate assembly protocol was used for part assembly and running the Golden Gate reaction in the OT2. A chemical transformation protocol relying on the temperature block was applied to transform the plasmids in the ST18 conjugation strains. A supplementary plating protocol was used as a flexible choice for variable plating options. These protocols are detailed in the Supporting Information 2.

## Results and discussion

### Domestication and characterization of parts

#### Assessment of origins of replication

Several origins of replication (ORIs) have previously been used in *R. sphaeroides* (Davis et al., 1988; Zeilstra-Ryalls & Kaplan, 1995), but to date, only the pBBR1 ORI has been utilized within a toolbox framework (Huo, 2011; Tikh et al., 2014). We characterized two additional broad-host ORIs – RK2 and RSF1010 – and incorporated them as standard conjugational entry plasmids with several antibiotic resistance genes in both level 1 and 2 to extend the Zymo-Parts toolbox. The ORIs have been added to the toolbox before, but only as single level 1 expression vectors with a limited range of antibiotic resistance genes (Behrendt et al., 2024; Wichmann et al., 2023). We assessed their functionality by constructing plasmids containing the respective ORI, an *egfp* expression cassette and a kanamycin resistance selection marker (Figure 2A). All three plasmids were successfully conjugated into *R. sphaeroides,* confirming the ability of all three ORIs to support plasmid maintenance and replication within the host. Quantitative fluorescence analysis revealed reproducible differences in expression levels among the ORIs. The RSF1010 ORI-based plasmid resulted in the highest biomass-normalized fluorescence intensity (from here on referred to as normalized fluorescence), 1.35-fold higher compared to the pBBR1-based plasmid (Figure 2B). The RK2 ORI-based plasmid led to a 1.2-fold higher normalized fluorescence compared the pBBR1-based plasmid. The differences in *egfp* expression levels between the three ORIs were relatively small, suggesting that the plasmids are present in similar copy numbers in *R. sphaeroides*.

**Figure 2.**
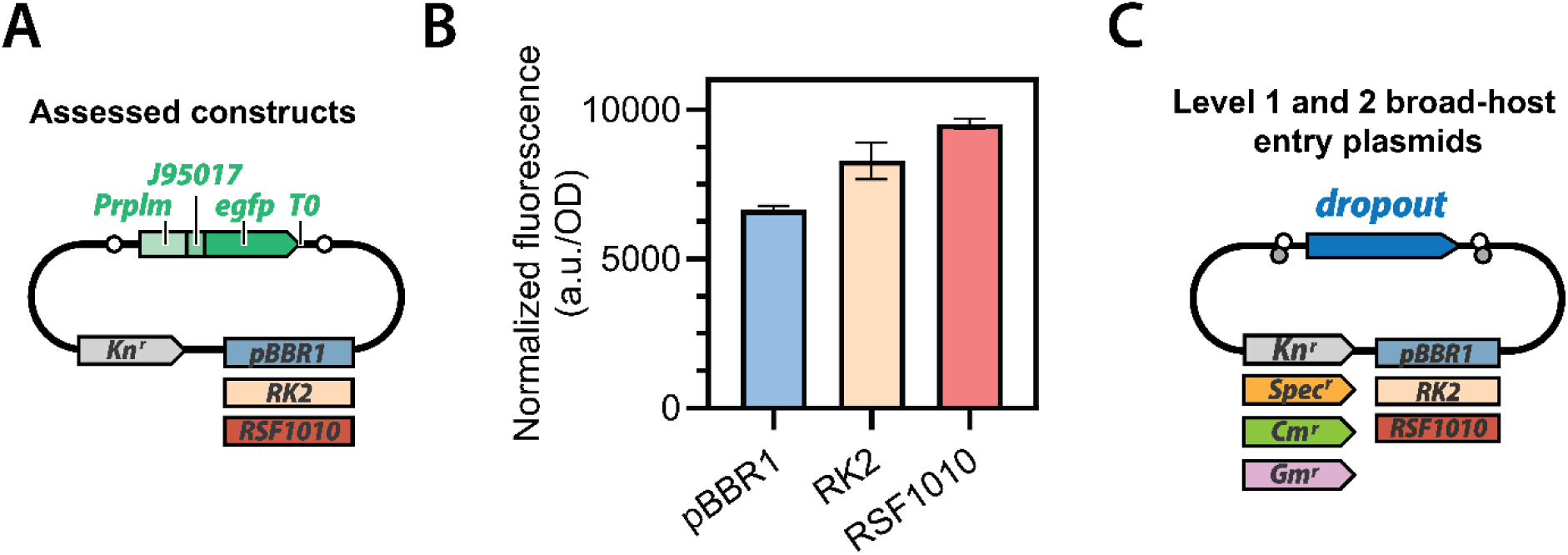
Characterization of the impact of origins of replication on egfp expression in R. sphaeroides. A) Schematic representation of the assessed constructs B) Biomass-normalized fluorescence intensity of exponentially growing R. sphaeroides containing egfp expression plasmids with different ORIs. Error bars represent the standard deviation of technical triplicates. C) Extension of the modular cloning toolbox to include broad-host entry vectors for both level 1 and 2 plasmids with different antibiotic resistance markers.

These findings align with previous assessments of the three ORIs in *E. coli*. Damalas *et al*. (2020) showed that pBBR1-based constructs yielded similar expression levels to the RK2-based constructs, while RSF1010-based constructs demonstrated ∼1.5-fold higher expression. The plasmid copy numbers quantified in *E. coli* (3-4 for pBBR1, 4-7 for RK2 and 7-10 for RSF1010) are consistent with these expression level observations (Durland et al., 1990; Lee et al., 2011; Meyer, 2009; Tao et al., 2005). Of note, in *P. putida,* pBBR1 and RSF1010-based plasmid expression resulted in 2-3 fold increased expression levels compared to the RK2-based expression (Damalas et al., 2020).

To extend the flexibility of the modular cloning system and enable orthogonal selection strategies, we constructed level 1 and level 2 entry plasmids with the three ORIs combined with resistance markers for four commonly used antibiotics: kanamycin, gentamicin, spectinomycin and chloramphenicol (Figure 2C, Supplementary Note 5). A list of all plasmids available in the toolbox is shown in Supplementary Table S2. Our expansion of the Zymo-Parts toolbox with a wide array of antibiotic resistance markers and broad-host ORIs enables an easy adoption of the toolbox to other non-conventional bacterial hosts.

#### Assessment of promoter strength

Gene expression can be modulated at the level of transcription initiation, where the promoter sequence plays an important role in determining transcriptional rates. To expand and characterize the collection of promoter sequences available for *R. sphaeroides,* we assessed the activity of three native promoters (P_J95023_, P_J95025_, and P_J95027_; Huo, 2011), two heterologous promoters originating from *Gluconobacter oxydans* and *Paracoccus zeaxanthinifaciens* (P*_rpmI_*, P*_crtE_*) and eight synthetic promoters (P_J23100_, P_J23104_, P_J23105_, P_J23114_, P_J23116_; Anderson, (2006)) (P_strong1k_, P_strong10k_, and P_strong100k_; Behrendt et al., 2022). The P_J23104_ promoter contained a point mutation and is referred to as P_J23104*._ The synthetic promoters sourced from the Anderson collection (2006) are widely adopted by the synthetic biology community and have been assessed in other organisms such as *E. coli, P. putida and Cupriavidus necator* (Damalas et al., 2020; Mishra et al., 2024; Stukenberg et al., 2021).

To assess promoter activity, we constructed plasmids with transcriptional units consisting of the tested promoter, RBS J61100, *egfp* and the T0 terminator, all integrated into the pBBR1 ORI-based backbone with a kanamycin resistance marker (Figure 3A). The *R. sphaeroides* strains carrying the respective plasmids were cultivated in 96-well plates with continuous monitoring. The normalized fluorescence of the strains carrying all constructs remained constant throughout the exponential growth phase (Figure S9). We observed a 270-fold dynamic range in normalized fluorescence between the strongest and weakest promoters (promoters P_J23100_ and P_J95023_, Figure 3A).

**Figure 3:**
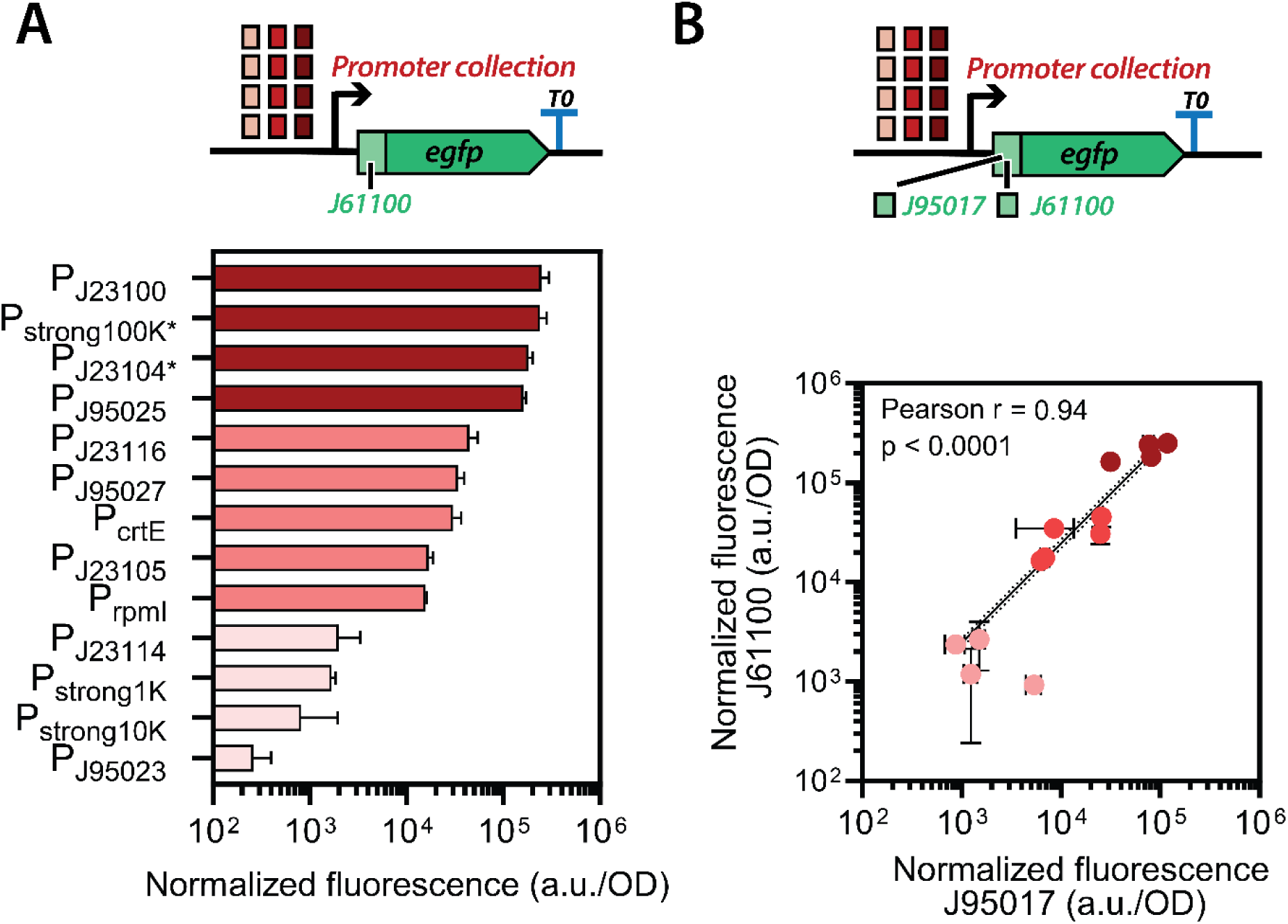
Assessment of promoters in R. sphaeroides. A) Ranking of the promoter collection based on the biomass-normalized fluorescence values. The promoters were assessed in combination with RBS J61100. Error bars represent the standard deviation of technical triplicates B) Correlation of promoter strengths in R. sphaeroides when combined with either RBS J61100 or J95017. Horizontal and vertical error bars represent the standard deviation of technical triplicates. The black line represents linear regression. Pearson r value was calculated using the Pearson correlation formula in GraphPad.

Strong synthetic promoters designed for *E. coli* were also the strongest in *R. sphaeroides,* with promoters P_J23100_, P_strong100k*_ and P_J23104*_. The strongest native promoter, P_J95025_, exhibited approximately 5-fold lower expression compared to the overall strongest promoter and, together with the synthetic P_J95027_, P_J23116_ and P_J23105_, and the heterologous P*_crtE_* and P*_rpmI_,* represent the medium strength collection of promoters. The remaining plasmids resulted in even lower normalized fluorescence, with P_J23114_ P_strong1k_, P_strong10k_, and P_J95023_ representing the lower end of the promoter collection.

Our results align with the work by Huo (2011), in which the promoters P_J95025_, P_J95023_ and P_J92027_ were designed and assessed. By benchmarking these previously established parts against the commonly used ones from the Anderson collection, we increased the available dynamic range by identifying promoters with both substantially higher and lower activities. Furthermore, the synthetic promoters showed a similar pattern in expression levels compared to the results obtained previously in *E. coli, V. natriegens* and *Z. mobilis* (Behrendt et al., 2022, Anderson, 2006, Stukenberg et al., 2021). The promoter P_J23104*_ resulted in 1.36-fold lower activity compared to P_J23100_, which is similar to the results obtained in *E. coli* by Anderson (2006) where the non-mutated P_J23104_ resulted in a 1.39-fold lower activity compared to P_J23100._ P_strong100k*_, P_strong10k_ and P_strong1k_ were designed by Behrendt *et al*. (2022) *in silico* using the DeNovoDNA promoter calculator (LaFleur et al., 2022). Since the elements appear to result in similar expression patterns in *R. sphaeroides*, more precisely tuned parts could be readily generated by such tools. This opens doors for the domestication of several other standard parts that are currently only used in model organisms. Furthermore, with this information, translation of genetic tools (Tn7 transposon-mediated recombination, CRISPRi, etc.) could become easier, not requiring extensive engineering efforts as demonstrated in the study by Hall *et al*. (2024).

All synthetic promoters based on the *E. coli* σ_70_ architecture were observed to also be active in *R. sphaeroides*, even though the promoter architecture in *R. sphaeroides* differs significantly from that of gammaproteobacteria such as *E. coli* (Myers et al., 2021). Our results support previous observations by Henry *et al*. (2020) and Hall *et al*. (2024) and show that the heterologous promoters with an orthogonal architecture can still allow for native sigma factor binding. Using such synthetic promoters may reduce native regulation by eliminating the dependence on the CarD transcription factor, which is required for most endogenous promoters (Myers et al., 2021).

After transcription, the promoter results in a short 5’UTR sequence that is still present in the mRNA and can influence the interaction between the ribosome and the Shine-Dalgarno sequence (Carrier & Keasling, 1997; Lou et al., 2012). To assess the modularity of promoter-RBS combinations and evaluate potential 5’UTR context effects, we compared expression profiles of the complete promoter collection when paired with either J95017 or J61100 RBSs (Figure 3B). The promoter collection combined with the J95017 RBS spanned a 135-fold range in normalized fluorescence, 2-fold less compared to when RBS J61100 (270-fold range) was used. The two datasets demonstrated a high level of correlation (Pearson r = 0.94, p < 0.0001; Figure 3B). Notably, P_J95023_ exhibited a 5.77-fold increase in normalized fluorescence, shifting from the weak group of promoters when coupled with RBS J6100 to the medium group with RBS J95017 (Figure S2). On the other hand, P_J95025_ and P_J95027_, the remaining native architecture-based promoters, both displayed the highest increase in normalized fluorescence, 5.15- and 4.09-fold, respectively. Other promoters resulted in a fairly consistent 1.5- to 3-fold decrease in normalized fluorescence when combined with RBS J95017 compared to RBS J61100 (Figure S1). The ranking of the promoter strengths remained consistent in the high and medium strength levels, with few inconsistencies observed in the weak range, where specific RBS-5’UTR interactions exhibited a more pronounced effect (Figure S2). Our data demonstrate that a change in the RBS coupled to the promoter collection results in predictable expression levels even in the absence of a dedicated insulator. However, if the insulator is desired for more advanced applications, an insulator RiboJ (a combination of a ribozyme and a hairpin, (Lou et al., 2012)) is available as a standard part of the Zymo-Parts MoClo toolbox (Wichmann et al., 2023).

#### Assessment of inducible expression systems

To expand the collection of inducible expression systems, we assessed four systems: (1) the salicylic acid inducible NahR-P*_salTTC_* (Meyer et al., 2019), (2), the vanillic acid inducible VanR-P*_vanCC_* (Meyer et al., 2019), (3) the IPTG inducible LacI-P_lacT7A1*_O3O4*_ (Deuschle et al., 1986) and (4) the 3-methyl benzoate (3-MB) inducible XylS-P*_m_* system (Mermod et al., 1986). References for these inducible expression systems can be further found in Schuster & Reisch (2021). We constructed plasmids containing transcriptional units with the respective inducible system (consisting of the regulatory element under the control of a constitutive P*_lacI_* promoter and the inducible promoter), together with the J61100 RBS, *egfp* and the T0 terminator, integrated in the pBBR1-based expression plasmid with a kanamycin resistance marker (Figure 4A). Growth and fluorescence of the conjugated *R. sphaeroides* strains were assessed in a plate reader with varying concentrations of inducers, allowing for the calculation of dose responses using the Hill equation.

**Figure 4:**
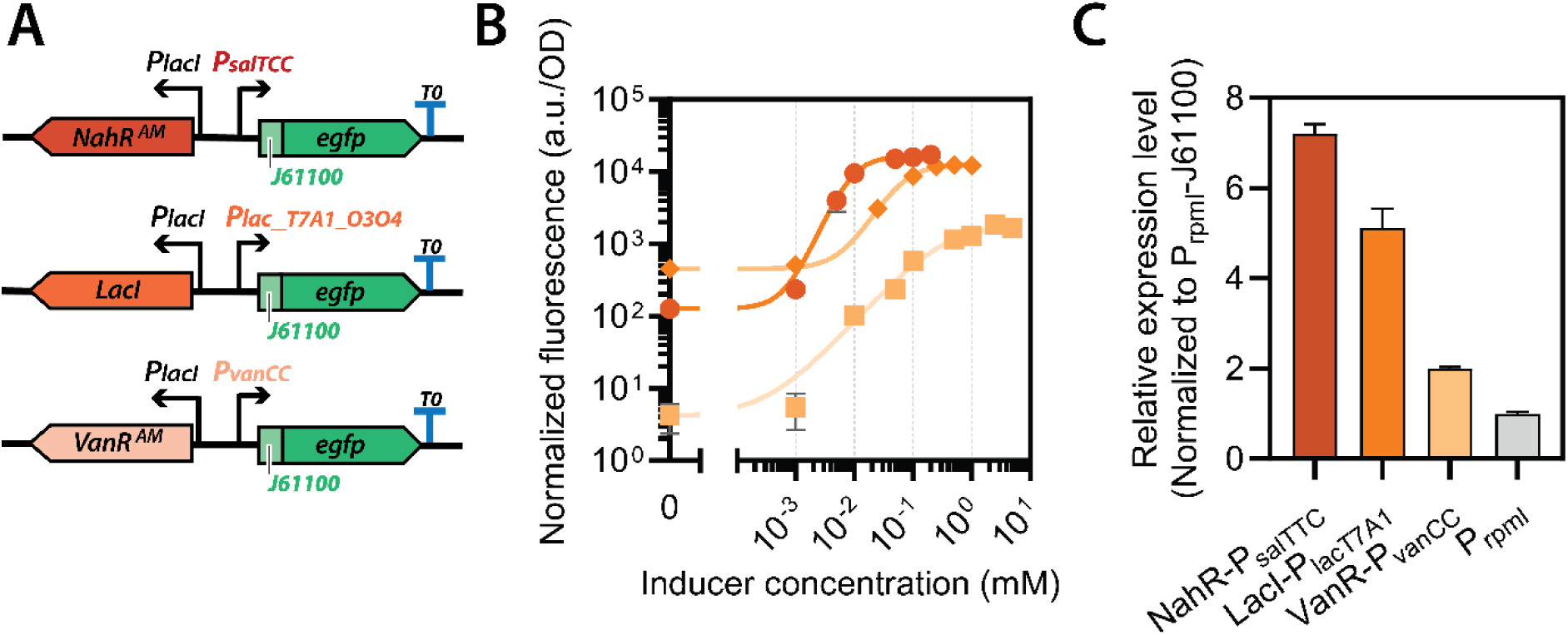
Characterization of inducible systems in R. sphaeroides. A) Schematic representation of the transcriptional units constructed for the assessment of inducible systems. All transcriptional units were assembled into the pZP1126 expression vector with pBBR1 ORI and kanamycin resistance marker. B) Dose-response curves of the inducible systems in the presence of varying concentrations of the inducers. The points represent the average normalized fluorescence at specific inducer concentration, while the line represents the fitted Hill equation (Methods). Color of both points and lines corresponds to the color of the bars in panel C. Error bars represent the standard deviation of technical triplicates. C) Comparison of expression levels of the inducible systems at maximal induction to a reference expression unit PrpmI_J61100). Error bars represent the standard deviation.

NahR-P*_salTTC_*, VanR-P*_vanCC_* and LacI-P*_lacT7A1_O3O4_* systems followed similar induction dynamics, reaching full induction within 4 h (Figure S3). The XylS-P_m_ system displayed a leaky expression, with a high level of basal normalized fluorescence in the absence of the inducer (Figure S4B). Furthermore, the inducer – 3-MB – negatively influenced growth at concentrations above 0.1 mM (Figure S4A). Due to the inducer’s toxicity and leaky expression, this system it is not a viable option for controlling gene expression in *R. sphaeroides*.

NahR-P*_salTTC_* and VanR-P*_vanCC_* systems exhibited 136-fold and 432-fold dynamic ranges of induction (fluorescence intensity ratio between the uninduced and maximally induced culture), respectively, while the LacI-P*_lacT7A1_O3O4_* system exhibited a 27-fold dynamic range (Figure 4B, Table 1). The exceptionally wide dynamic range observed in VanR-P*_vanCC_* is largely due to a very low level of basal expression of the system in uninduced conditions (Figure S3B). NahR-P_salTTC_ and LacI-P*_lacT7A1_O3O4_* exhibited a higher basal *egfp* expression when not induced; but still 20-fold and 5-fold lower compared to the P*_rpmI_*-J61100 expression, respectively (Table 1). However, when induced maximally the systems resulted in 7.2 and 5-fold higher expression levels compared to the medium strength promoter P*_rpmI_* (Figure 4C). The VanR-P*_vanCC_* demonstrated a lower expression level, albeit still 2-fold higher when compared to the P*_rpmI_*-J61100 (Figure 4C).

**Table 1:** Inducible systems assessed in R. sphaeroides.

|  | LacI-P<sub>lacT7A1_O3O4</sub> | NahR-P<sub>salTTC</sub> | VanR-P<sub>vanCC</sub> |
| --- | --- | --- | --- |
| Hill slope | 1.55 ± 0.22 | 1.89 ± 0.22 | 0.91 ± 0.1 |
| EC<sub>50</sub> [μM] | 59 ± 7 | 8.8 ± 0.5 | 310 ± 60 |
| Dynamic range [fold]<sup>a</sup> | 27 ± 2 | 136 ± 5.6 | 432 ± 192 |
| Basal expression level<br>[relative to P<sub>rpmI</sub>] | 0.19 ± 0.01 | 0.05 ± 0.004 | 0.005 ± 0.002 |

<sup>a</sup> Dynamic range is represented as the ratio between biomass-normalized expression levels between maximal induction and the basal expression in uninduced cultures.

While IPTG-based inducible systems have been described in *R. sphaeroides* before (Hall et al., 2024; Ind et al., 2009; Kretz et al., 2023), the use of the vanillic acid and salicylic acid-based systems is novel. The use of the latter as an inducer provides a cheaper alternative to the more expensive IPTG, facilitating bioproduction at industrial scales (Ferreira et al., 2018). . The expanded selection of inducible systems in this molecular toolbox allows dynamic regulation of multiple modules simultaneously, promising for both metabolic engineering and synthetic biology applications, including extensive genetic circuits or logic gates.

#### Assessment of RBS strength

In addition to different promoters, protein levels can also be modulated by changing the RBS, which influences the rate of translation. We assessed five synthetic RBSs designed for *E. coli* (B0030, B0031, B0032, B0034; (Mahajan et al., 2003)) (J61100; Anderson, 2007) and six designed for *R. sphaeroides* (J95015, J95016, J95017, J95018, J95019 and J95028; Huo, 2011) (Figure 5A). We constructed plasmids with transcriptional units consisting of the P*_rpmI_* promoter, the tested RBS, *egfp* and the T0 terminator, all integrated into the pBBR1 ORI-based backbone with a kanamycin resistance marker (Figure 5A). The *R. sphaeroides* strains carrying the assessed plasmids were grown in a plate reader where growth and eGFP fluorescence were measured.

**Figure 5:**
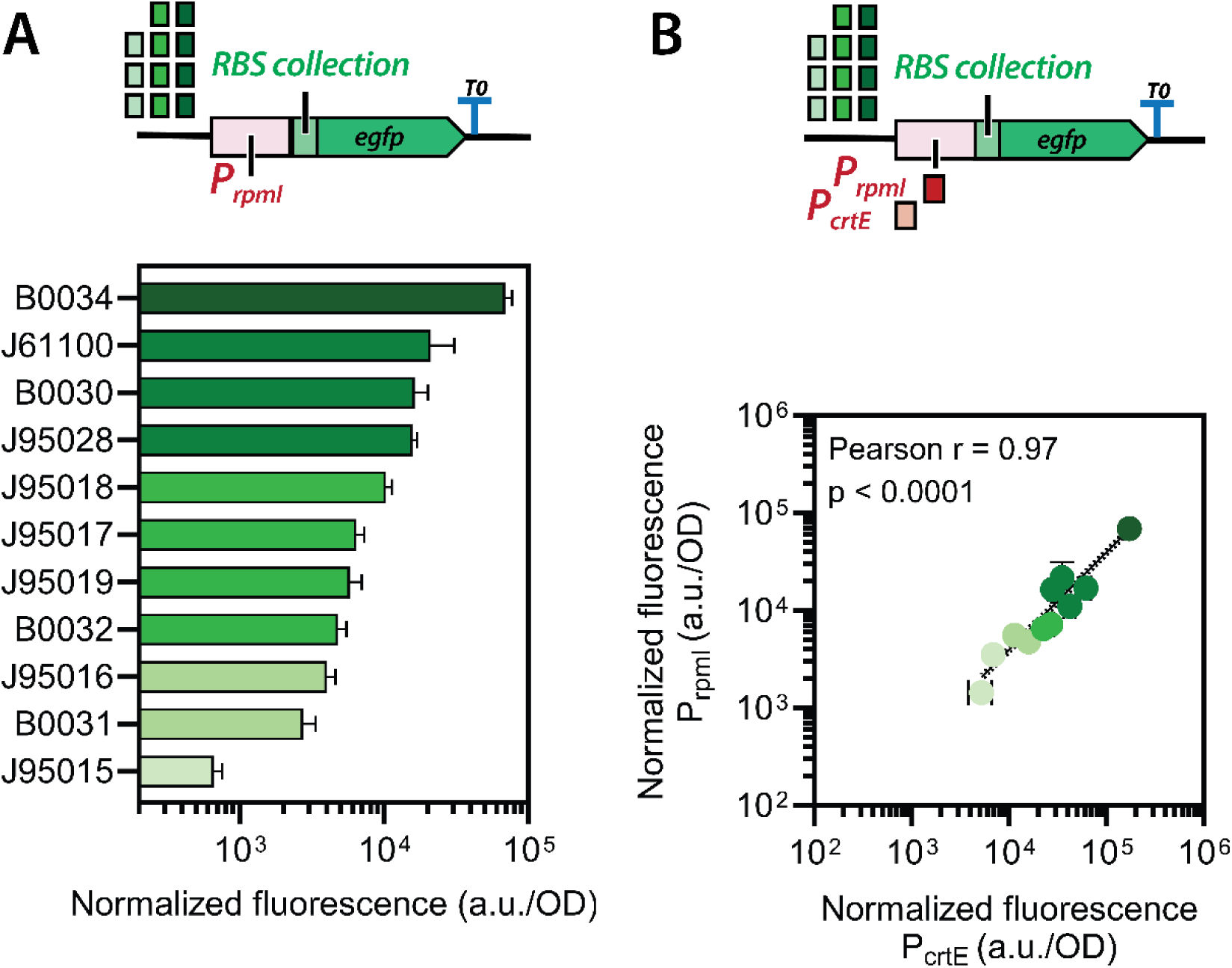
Characterization of RBSs in R. sphaeroides. A) Ranking of the RBS strengths when coupled with promoter P_rpmI_. Error bars represent standard deviation of technical triplicates. B) Correlation of the RBS strengths when coupled with either P_rpmI_ or P_crtE_. The vertical and horizontal error bars represent standard deviations of technical triplicates when coupled to the respective promoters. The black line represents linear regression. Pearson r value was calculated using the Pearson correlation formula in GraphPad.

A 49-fold dynamic range was observed among the tested RBSs, with B0034 and B0030 being 3.5- and 2-fold stronger compared to the strongest RBSs designed for *R. sphaeroides* (Figure 5A). Strengths of RBSs designed for *R. sphaeroides* are in good correlation to the results by Huo (2011), with the exceptions of J95015 and J95017. In the work of Huo *et al*., RBSs contained a 6-bp cloning scar as a consequence of BioBrick cloning, which acts as a spacer between the Shine-Dalgarno (SD) sequence and the start codon. This scar was omitted in our designs due to the implementation of Golden Gate cloning, resulting in only a single base-pair scar between the RBS and the start codon (Figure S5B). RBS J95015 displayed a lower activity compared to the results obtained by Huo (2011), which could be due to the SD sequence being present at the very 3’ end of the RBS, with only a single base spacing the element from the start codon (Figure S5). The addition of a standardized spacer to those parts might further increase expression levels. Nonetheless, even in the absence of this spacer, the majority of the RBSs resulted in normalized fluorescence comparable to that of medium-range *E. coli-*based RBSs, showing that the ribosome is flexible enough to still initiate translation.

Synthetic RBSs based on the native design led to lower expression levels compared to the *E. coli-*based designs. This is especially interesting since *R. sphaeroides’* 16S rRNA recognizes a different Shine-Dalgarno (SD) sequence (alignment of 16S rRNA sequences shown in Figure S6). The high compatibility of the *E. coli-*based RBSs opens opportunities to source these genetic elements from standard repositories, such as the iGEM registry of standard genetic parts (https://parts.igem.org/Main_Page).

We assessed our complete RBS collection under the control of promoters P*_rpmI_* and P*_crtE_* (Figure 5B). The dynamic range of the RBSs paired with P*_crtE_* spanned a 33-fold range, 1.5- fold lower compared to when paired with P*_rpmI_*. However, the reduction in the dynamic range is largely due to a change in expression of the weakest RBS, J95015, with the other RBS showing a more predictable response (Figure 5B), with a 1.7- to 4-fold increase compared to when combined with the P*_rpmI_* promoter (Figure S7). The expression levels of the RBSs when controlled by either P*_rpmI_* or P*crtE* promoter show a high correlation (Pearson r = 0.97, p < 0.001) (Figure 5B). The RBSs act predictably and retain an ordered hierarchy, with the utilized promoter not influencing the final RBS strength ranking (Pearson r = 0.95, p < 0.0001; Figure S8). Our findings align well with a recent study performed in *Staphylococcus aureus* by Rondthaler *et al*. (2023): when comparing non-insulated RBSs with two promoters, they also observed a good correlation for most of the assessed parts.

Overall, we show that the collection of *R. sphaeroides* RBSs is robust and spans an up to 104-fold dynamic range, depending on the promoter used. Together with the polycistronic operon-building capabilities of the toolbox, this RBS collection opens possibilities to precisely control gene expression within single transcriptional units that are under the transcriptional control of the same promoter. The characterization also demonstrates the opportunity for further optimization of RBS designs, and further research should focus on incorporating standard spacers into the existing parts.

### Combinatorial assembly guided characterization of genetic elements

#### Integration of Rhodo-box into a semi-automated workflow

To further study the combined effect of various promoter and RBS sequences on expression strength, we developed a semi-automated workflow to cross-screen a combinatorial library of parts. Traditional genetic part characterization is typically performed by comparing different elements of the same class (e.g., promoters) within a fixed genetic context (RBS, CDS, terminator) (Bak et al., 2022; Englund et al., 2016). While this approach provides valuable baseline performance data, its translatability to diverse genetic contexts is often unpredictable due to context-dependent effects including mRNA secondary structure formation, transcriptional interference, and regulatory element interactions (Köbbing et al., 2020).

To systematically assess context-dependent interactions and validate the modular design principles underlying the Rhodo-Box toolkit, we leveraged a semi-automated workflow to construct and characterize a comprehensive combinatorial library of genetic circuits. Using this combinatorial approach, we assembled 5 promoters with 6 RBSs, and 4 terminators with two pairs of promoters and RBSs, with an overall success rate of 90%. Ultimately, 38 strains were built within 5 days in a single workflow (Figure 6A). In contrast to the estimated 4 h of manual handling required for this workflow, the semi-automated approach reduced the intervention time to 2 h, reducing by 50 % the allocated time required to build this strain library (see Supporting Information 2).

**Figure 6.**
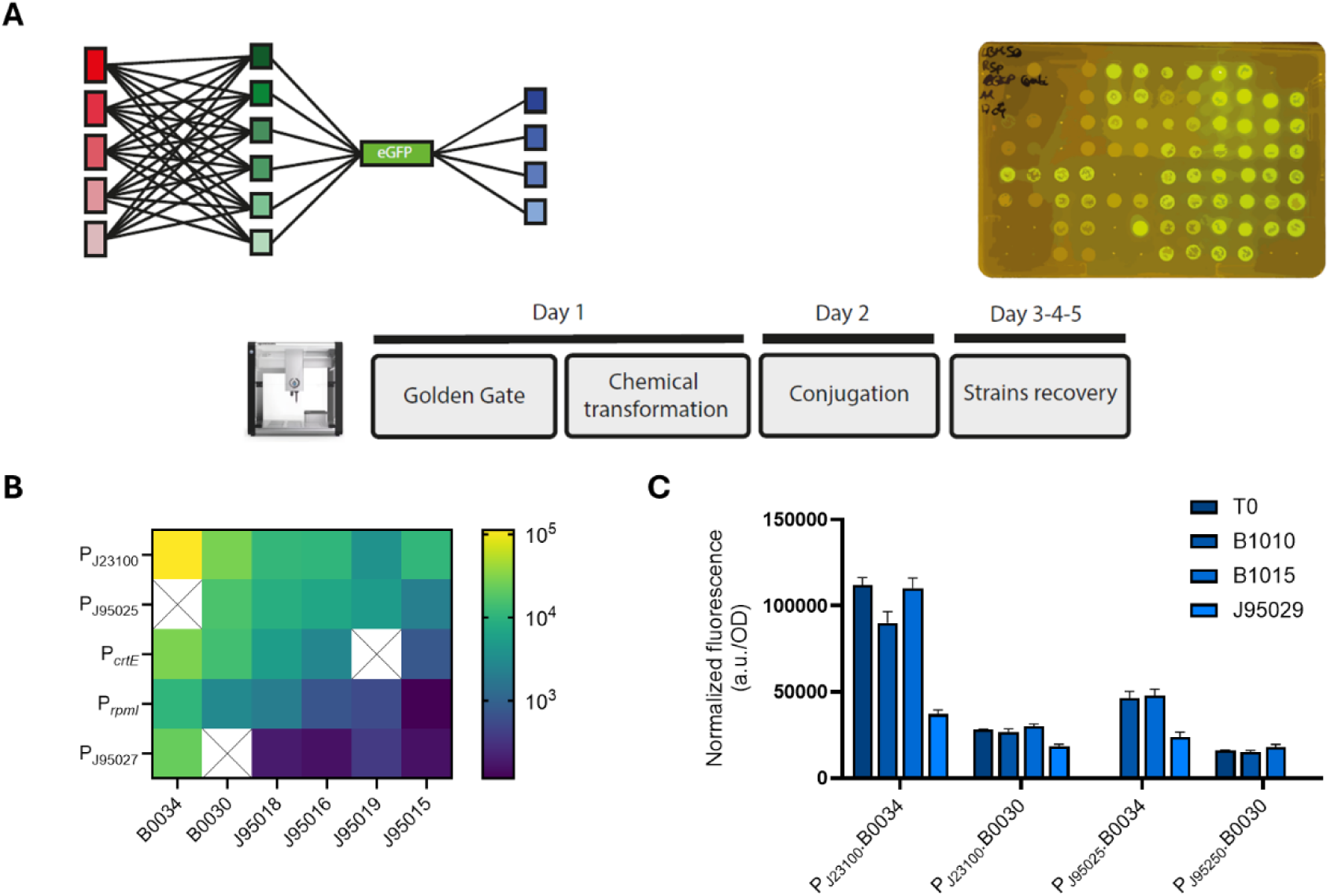
Combinatorial part characterization using a semi-automated cloning workflow. **A)** Semi-automated workflow, combining 5 promoters (red) with 6 RBSs (green), and characterizing 4 terminators (blue) with an egfp. Using the Opentrons OT2, only five days are required to build a level 1 R. sphaeroides strain library from a level 0 plasmid library. **B)** Heat-map representing the combinatorial promoters and RBSs strengths in R. sphaeroides. **C)** Normalized fluorescence of terminators T0, B1010, B1015, J95029 expressed in combinations of P_J23100_ and P_J95025_ with B0034 and B0030. Graphs B) and C) represent the average fluorescence values normalized with OD_600_, the standard deviation of three technical replicates is represented in C).

#### Assessment of combinatorial promoters and RBSs interactions

Five promoters from various origins were selected among the strong and intermediate strengths for this combinatorial assembly. The synthetic promoter P_J23100_ consistently achieved the highest expression levels across all RBS combinations, confirming its robust performance as the strongest element in our collection (Figure 6B). The only exception occurred in combination with RBS J95019, where P_J23100_ and P_J95025_ promoters achieved comparable expression levels, likely reflecting the limiting effect of inefficient ribosome binding under these conditions. Among the medium-strength promoters, distinct performance patterns emerged. P_J95025_ was stronger than the P*_crtE_* promoter in most contexts, except when paired with RBS B0030, where it demonstrated comparable expression levels. The P*_crtE_* promoter was consistently about 4-fold stronger than P*_rpmI_*. This is in line with our previous observation in the “Assessment of RBS strength” section.

In this combinatorial promoter comparison, three promoters (P_J95025_, P*_crtE_* and P*_rpmI_*) demonstrated a proportional decrease in strength when paired with successively weaker RBSs, indicating predictable relationships that support modular design principles. RBS strength hierarchy and proportionality appear to be conserved across these promoters, despite their different origins. In contrast, P_J23100_ and P_J95027_ shared a similar pattern of fixed strengths when paired with the synthetic *R. sphaeroides* RBSs (J95015, J95016 and J95018). These promoters might interact with the specific sequence of these RBSs derived from the same SD sequence. Despite using promoters from different sources, proportional promoter strengths seem to be conserved among most RBSs (Figure S15A). A maximum 91-fold dynamic range was measured for the promoters in this combinatorial approach, providing substantial flexibility for expression tuning in metabolic engineering applications.

Six RBSs were selected in this combinatorial study to cover the dynamic range observed in the previous characterization in the “Assessment of RBS strength” section. When combined with various promoters, RBS B0034 was consistently the strongest across all combinations, while B0030 was the second strongest RBS (Figure 6B). Among the synthetic *R. sphaeroides* RBSs, J95018 outperformed J95016 with P_J95025_ (1.3-fold), P*_crtE_* (1.9-fold), and P_rpmI_ (2.9-fold), but showed similar strength when paired with the P_J23100_. RBS J95016 was stronger than J95019 with P_J23100_ (2.7-fold), P_J95025_ (1.4-fold) and P_rpmI_ (1.3-fold). Finally, J95015 was systematically the weakest RBS, except with P_J23100_ and P_J95027_ where it achieved similar strength to J95018 and J95016. In this combinatorial comparison of RBSs, J95018 and J95016 have a conserved proportionality and hierarchy between promoters. However, this behavior is not observed with RBSs J95019 and J95015. Even though the synthetic *R. sphaeroides* RBSs are derived from the same source, this did not seem to guarantee a consistent strength expression between all promoters (Figure S15B). A maximum 146-fold dynamic range was measured for the RBSs in this combinatorial approach, highlighting the RBSs ability to largely increase the dynamic range compared to solely varying promoters, enabling precise fine-tuning of expression strength in transcriptional units.

In this combinatorial parts characterization, we observed a 961-fold dynamic range in expression strength. It is worth noticing that the weakest promoters from our library were not included in this combinatorial characterization, potentially offering an even wider dynamic range for the overall library. Despite a general conservation of hierarchy and proportionality between promoters and RBSs, occasional inconsistencies can be observed, likely due to the impact of the specific genetic context. In a metabolic engineering setting, this dependency can also be observed with the 5’-end of alternative coding sequences (Kosuri et al., 2013). Therefore, the construction of combinatorial libraries for metabolic engineering applications enables an increased throughput to screen large pools of strains, allowing to overcome unpredicted suboptimal designs.

#### Assessment of context-dependent terminators impact

Transcriptional terminator collections are usually limited in most toolbox part libraries. The use of the same DNA sequence when assembling multiple transcriptional units in level 2 plasmids could lead to undesired homologous recombination, especially in case of expression of burdening cargos. In this study, we compared the impact of four terminators on the expression levels of their upstream transcriptional unit. These four terminators were studied in a combinatorial approach: T0 terminator from phage lambda, B1010 and B1015 artificial terminators from Huang (2007) and a BioBrickTM bidirectional terminator-Omega terminator J95029 from Huo (2011). When combined with the P_J23100_-B0034-*egfp* construct, terminators T0 and B1015 showed similar fluorescence levels (Figure 6C). In contrast, B1010 led to slightly lower fluorescence, corresponding to 82% of the intensity observed with T0 and B1015, while J95029 resulted in a pronounced decrease (32% of T0/B1015). For the P_J23100_-B0030-*egfp* construct, T0 and B1010 gave comparable fluorescence (90% of B1015), whereas J95029 again reduced fluorescence significantly (61% of B1015). In the P_J95025_-B0034-*egfp* background, B1010 and B1015 produced equivalent fluorescence, while J95029 reduced expression by 51%. Finally, for P_J95025_-B0030-*eGFP*, T0 and B1010 yielded similar fluorescence, with B1015 slightly higher by 14%. Overall, with RBS B0030, terminator B1015 consistently gave marginally higher fluorescence compared to T0 and B1010 for the two tested promoters. The characterized terminators had a limited impact on the final expression strength of the transcriptional unit, except for J95029 with an average 2-fold strength decrease.

## Conclusions

Our work lays the foundation for a standardized Modular Cloning toolbox in *R. sphaeroides*, by greatly extending the *Zymo-Parts* toolbox, which has so far only been used for *Z. mobilis* and *E. coli*. We characterized 32 genetic parts for their use in *R. sphaeroides*, showing their robustness and wide dynamic ranges (270-fold for the promoter collection and 49-fold for the RBS collection), thus allowing for precise selection of parts depending on the final engineering goal. For the first time we show that three inducible expression systems in *R. sphaeroides* (NahR-P*_salTTC_*, VanR-P*_vanCC_* and LacI-P_lacT7A1*_O3O4*_) enable tight and dynamic (135-fold, 432-fold and 27-fold, respectively) expression control. We characterized and validated the functionality of three broad-host ORIs, which we included in the extended Zymo-Parts modular cloning toolbox as level 1 and 2 entry vectors, each combined with one of the four commonly used antibiotic resistance markers (kanamycin, gentamycin, spectinomycin, and chloramphenicol). Finally, we showcased the accessibility of a semi-automated cloning workflow for MoClo combinatorial assemblies, enabling fast and efficient building of *R. sphaeroides* libraries.

## Author contributions

**M.K**. and **A.R.** conceived the study, designed the experiments, analyzed the data, and wrote the manuscript. **G.D.** performed the characterization of the promoters, RBSs, and the ORIs. **M.K.** performed the characterization of the inducible systems. **A.R.** performed the automation study. **G.B.** contributed to the conceptualization and writing of the study. **V.A.P.M.d.S., J.B., C.B., R.A.W., E.A.G.**, and **M.M.M.B.** supervised the research and contributed to writing and reviewing of the study.

## Acknowledgements

This work was developed in the context of the BIOS project and has received funding from the European Union’s Horizon Europe research and innovation program under the HORIZON-RIA action, grant agreement No. 101070281.

The work was also supported by Isobionics B.V.

## Additional information

Supporting information file 1: Contains supporting figures, tables, and notes.

Supporting information file 2: Contains automation protocols

Sequences of the part collection are available at Zenodo: 10.5281/zenodo.17493661

