## Supporting Information for "Rhodo-Box: a Synthetic Biology Toolbox to Facilitate Metabolic Engineering of *Rhodobacter sphaeroides*"

### Supplementary notes

#### Supplementary note 1: Part domestication within Level 0

For propagation of basic entry vectors (level 0), a ColE1 origin pUC19-based plasmid was used, as described by Behrendt et al. (2022). The entry position of entry vectors was determined by specific four base-pair overhangs.

To introduce novel parts into the level 0 collection, the DNA provided was either as a hybridized double-stranded oligonucleotide (IDT), a gene fragment (Twist) or a PCR product. In any case, the sequence of interest was flanked by BbsI recognitions site and a four-base sequence complementary to the overhangs of the entry positions. Promoter sequences were assembled into vector pZP001, RBS sequences into pZP002, coding sequences into pZP003 and terminator sequences into pZP004.

For entry parts shorter than 100 bp (including flanking BbsI recognition sites), the DNA was ordered as two complementary single stranded oligonucleotides (IDT). The oligonucleotides (complete list in Table S4) were resuspended to 1 µM in Elution buffer (Thermo Scientific) and annealed using a temperature drop protocol (30 s 95 °C, 30 s 90 °C, 30 s 85 °C, 30 s 80 °C, 30 s 75 °C, 30 s 70 °C, 30 s 65 °C) in a thermocycler. 50 fmol of the resulting double stranded oligonucleotide was added to the Golden Gate assembly reaction. To domesticate DNA sequences from other, already existing vectors, a PCR using Q5 high-fidelity polymerase was performed using primers that added the BbsI recognition sites with specific four-base overhangs. If present, internal BbsI and BsaI recognition sites within the coding sequence were removed via the domestication PCRs and subsequent assembly of multiple parts. Following gel electrophoresis, the fragments were cut from the gel and purified using GeneJet Gel Extraction and DNA Cleanup Micro kit (Thermo Scientific) before being used in the Golden Gate assembly reaction.

To assemble parts into level 0 vectors, briefly, 1 µL of the 10X CutSmart buffer (NEB), 0.5 µL of recombinant bovine serum albumin (rBSA, NEB), 0.5 µL of the T4 ligase (NEB) and 0.5 µL of the BbsI restriction enzyme (NEB) were mixed with 1 µL of the required entry vector and 0.5 µL of the toolkit part. The volume was adjusted to 10 µL using MQ $H_2O$. For the reaction, 30 cycles of 3 min at 37 °C, followed by 5 min at 16 °C were carried out, and ended with a final digestion of 10 min at 37 °C, and a final inactivation of 10 min at 80 °C.

#### Supplementary note 2: Construction of polycistronic operons

The operon fragments can be constructed by combining an RBS with a GOI with dedicated level -1 entry vectors. The entry vectors carry specific four-bp overhangs for subsequent BbsI assembly into level 0. These entry vectors determine the position of the RBS-GOI operon fragments within the overall polycistronic operon. Currently, the toolbox supports up to five RBS-GOI within an operon unit, but extension is possible. Each position also contains a separate entry vector with the right overhang matching the entry vector of the subsequent level 0 plasmid, thus allowing for construction of custom-length operons. The desired complementary operon fragments can then be assembled into a standard level 0 polycistronic operon (position 7, spanning RBS and promoter position). A transcriptional level 1 unit can then be assembled within an expression vector by adding a promoter and a terminator, as described in Supplementary note 3.

#### Supplementary note 3: Construction of Level 1 expression vectors

Level 1 plasmids were assembled as described by Behrendt *et al.* (2022). Briefly, 1 µL of the 10X CutSmart buffer (NEB), 0.5 µL of recombinant bovine serum albumin (rBSA, NEB), 0.5 µL of the T4 ligase (NEB) and 0.5 µL of the BsaI-HF restriction enzyme (NEB) were mixed with 1 µL of the required entry vector and 0.5 µL of the toolkit parts. The volume was adjusted to 10 µL using MQ $H_2O$. For the reaction, 30 cycles of 3 min at 37 °C, followed by 5 min at 16 °C were carried out, and ended with a final digestion of 10 min at 37 °C, and a final inactivation of 10 min at 80 °C.

#### Supplementary note 4: Utilization of a double-loop limitless assembly

Level 0 parts can also be assembled into positioned level 1 vectors using BsaI assembly. The entry level 1 vectors in respective positions provide BbsI restriction sites and contain adjacent complementary overhangs, allowing for further integration into positioned level 2 vectors by BbsI assembly. Likewise, the entry level 2 vectors provide BsaI restriction sites and adjacent complementary overhangs, enabling double loop assembly “back” into level 1. The restriction enzyme for subsequent assembly into higher or lower level is added by the entry vector, meaning that the system is recursive and infinite, always adding and removing the respective restriction sites.

#### Supplementary note 5: Construction of broad-host entry vectors

Broad-host entry vectors were constructed using Golden Gate cloning. To construct additional Level 1 position 4 entry vectors, the backbones RK2 (pZP1125), pBBR1 (pZP1126), ORIs RSF1010 (pZP1395) were amplified with Q5 PCR using primer pair PR462 & PR463. The primers added SapI recognition sites and 3-bp overhangs which were compatible with the subsequently amplified antibiotic cassettes. The antibiotic cassettes were amplified using Q5 PCR from plasmids pSEVAb63-EV (GmR; Silva-Rocha et al., 2013), pZP334 (SpecR; Behrendt et al., 2022), and pZP331 (CmR, Behrendt et al., 2022) using primer pairs that added the SapI recognition sites and complementary 3-bp overhangs – PR460 & 461, PR458 & 459, and PR 456 & PR457, respectively. The PCR products were purified and assembled in a Golden Gate reaction using the SapI (NEB) restriction enzyme.

Level 2 position 1 entry vectors were constructed by amplifying each newly constructed Level 1 position 4 entry vector with specific primer pairs: **a)** pZ1W-E001, pZ1W-E002, and pZ1W-E003 with PR466 & PR467, **b)** pZ1W-E004, pZ1W-E005, and pZ1W-E006 with PR468 & PR470, and **c)** pZ1W-E007, pZ1W-E008, and pZ1W-E009 with PR468 & PR 469. The primers introduced a SapI recognition site with a defined 3-bp overhang. The mCherry vector was amplified from a plasmid pZP597 using primer pair PR464 & PR 465. The primers introduced level 2 position 1 determining BsaI and BbsI recognition sites and adjacent sequences and a SapI recognition site and a 3-bp sequence complementary to the one introduced to the pZ1W-E backbones. The PCR products were purified and assembled in a Golden Gate reaction using the SapI (NEB) restriction enzyme.

### Supplementary Figures

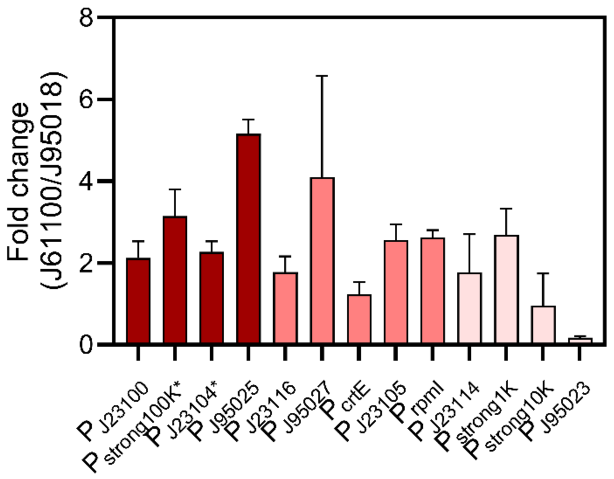


*Figure S1: Biomass-normalized fluorescence fold-change values of the promoter collection combined with either J61100 or J95018 RBSs. The error bars represent the standard deviation.*

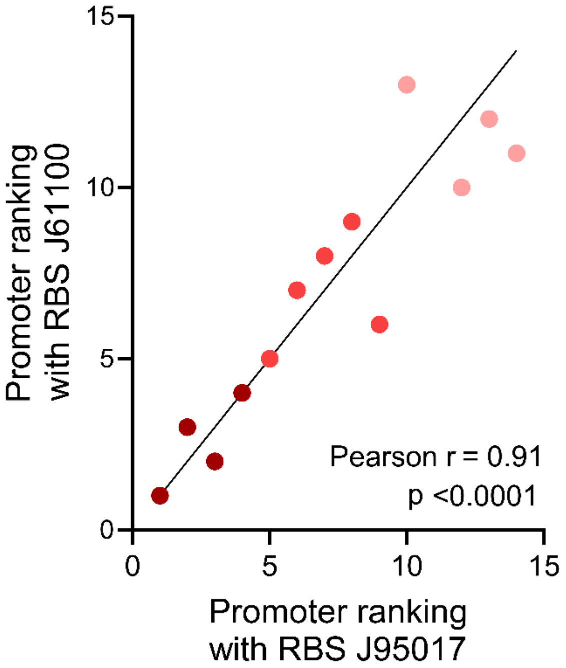


*Figure S2: Promoter strength ranking when coupled to either RBS J61100 or J95017, based on data shown in Figure 3. The black line represents a straight line with a hypothetical 1:1 relationship for reference. Statistical analysis was performed using GraphPad prism.*

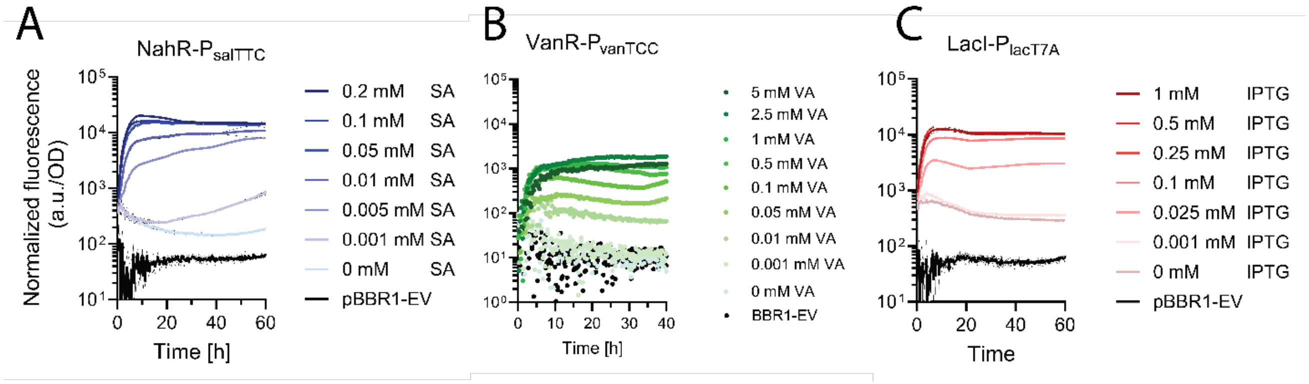


*Figure S3: Characterization of inducible systems. The points represent biomass-normalized fluorescence values. A) Assessment of the NahR-P$_{salTTC}$* system with different salicylic acid (SA) concentrations. B) Assessment of the VanR-P$_{vanTCC}$ system with different vanillic acid (VA) concentrations. C) assessment of the LacI-P$_{lacT7A}$ system with different IPTG concentrations. *Lines represents averages of a technical triplicates.*

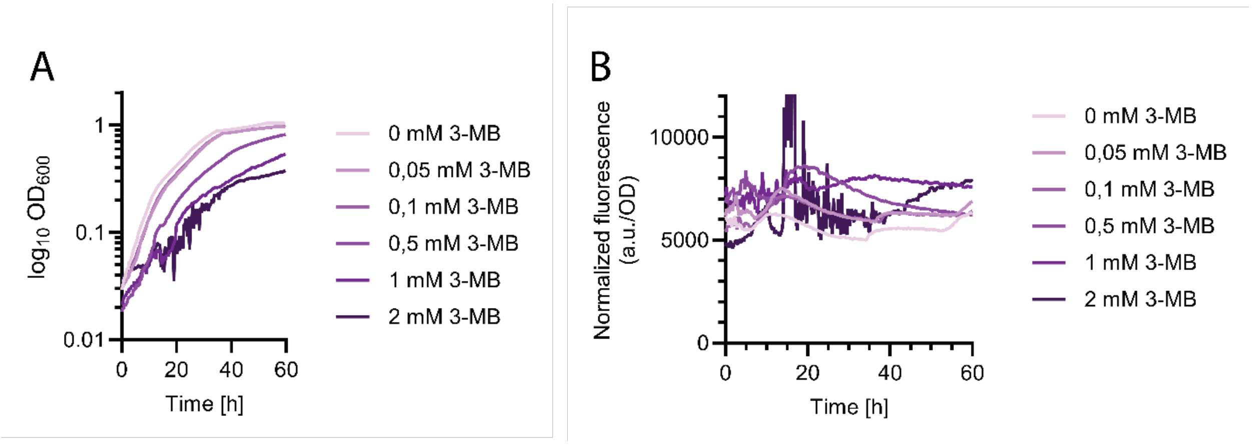


*Figure S4: Characterization of the XylS-Pm inducible system. A) Growth of the strain expressing the plasmid with the inducible system, J61100, egfp, T0 with pBBR1 ori and KnR marker. The line represents the average of a biological triplicate. B) Biomass-normalized fluorescence with different 3-MB concentrations. Each line represents the average of a technical triplicate.*

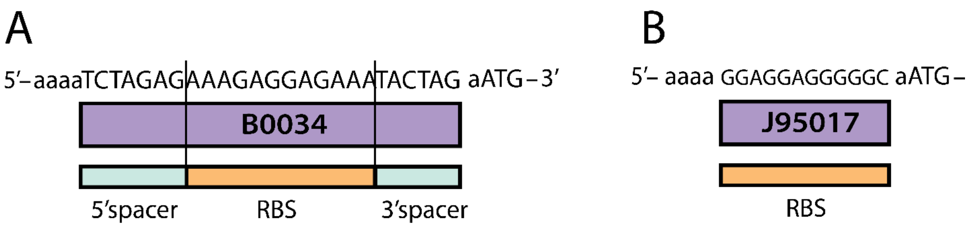


*Figure S1: Comparison of RBSs designed for A) E. coli, B) R. sphaeroides. Note that only B0034 contains the spacer sequence before the start codon ATG.*

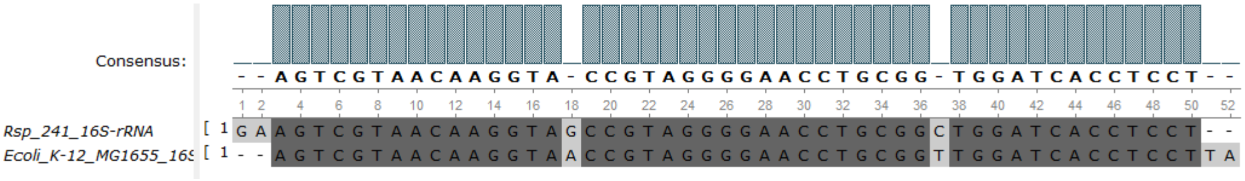


*Figure S2: Alignment of the last 50 bp of 16S rRNA sequence from R. sphaeroides 2.4.1 genome, and the E. coli K-12 subst. MG1655. The sequences were aligned using MAFFT. The agreements in the alignment are shown in dark grey color, while the differences are shown in light grey. Note that the 3' anti-SD sequence is different in the two organisms (addition of TA bases in the E. coli sequence).*

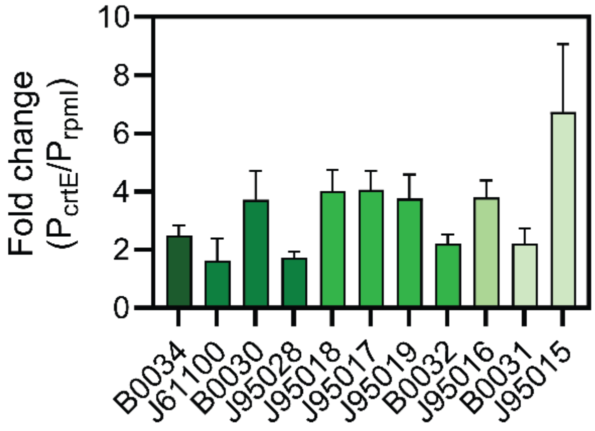


*Figure S7: Biomass-normalized fluorescence fold-change values of the RBS collection combined with either $P_{crtE}$ or $P_{rpmI}$ promoters. Error bars represent the standard deviation.*

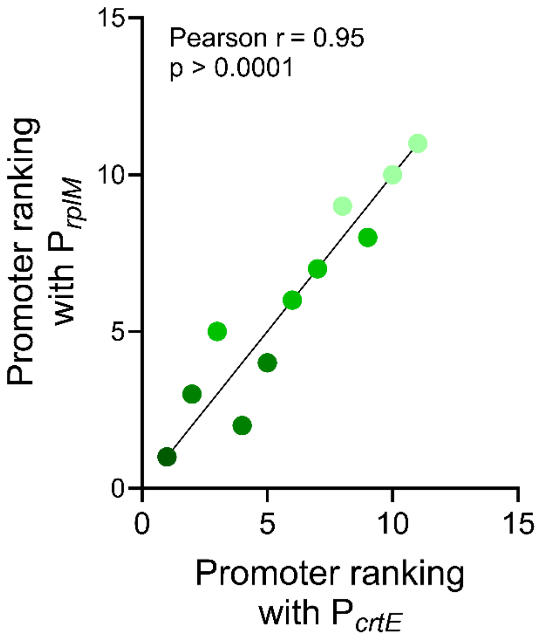


*Figure S8: RBS strength ranking when coupled to either $P_{rpmI}$ or $Pc_{rtE}$, based on data shown in Figure 5. The black line represents a straight line with a hypothetical 1:1 relationship for reference. Statistical analysis was performed using GraphPad prism.*

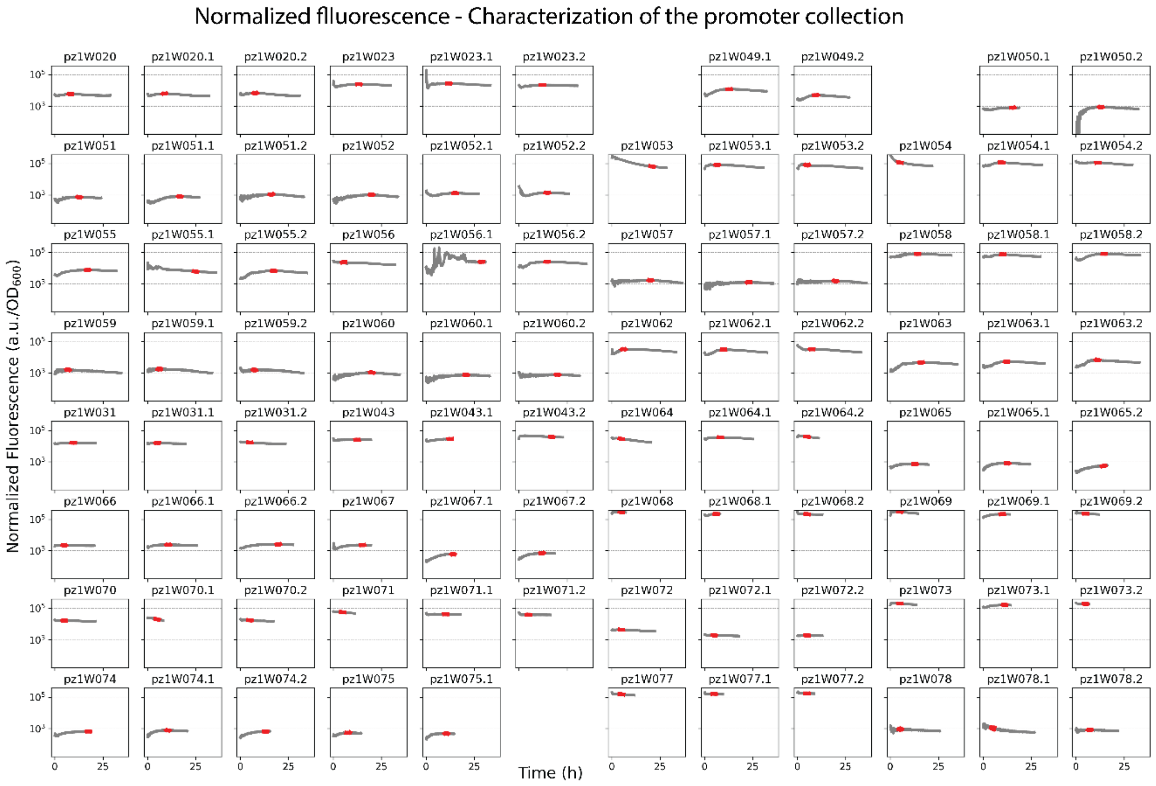


*Figure S3: Biomass-normalized fluorescence values of the promoter assessment. Each plot represents a well in a 96-well plate reader. The highest moving average of 25 points (excluding initial 4 in all samples except for wells C5, B7, C2, and E6 – 24, 20, 24, and 16 hours were excluded to obtain a stable average, respectively) used for the calculation of the average biomass-normalized fluorescence intensities is highlighted in red.*

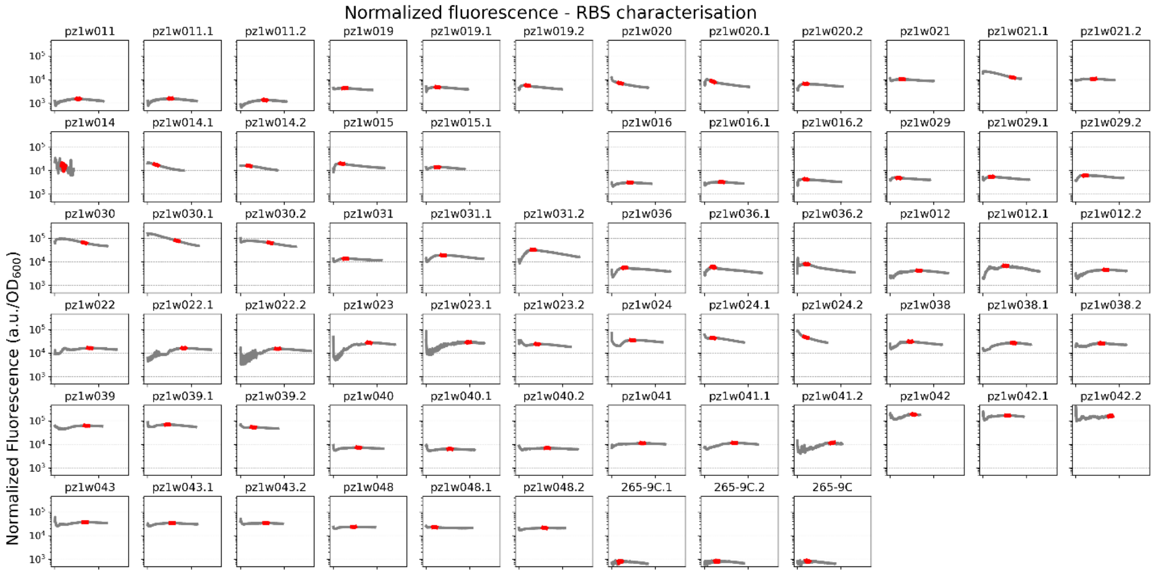


*Figure S4: Biomass-normalized fluorescence values of the RBS assessment. Each plot represents a well in a 96-well plate reader. The highest moving average of 25 points (excluding initial 4 in all samples except for wells C5 and B7 – and 24 and 20 hours were excluded to obtain a stable average, respectively) used for the calculation of the average biomass-normalized fluorescence intensities is highlighted in red.*

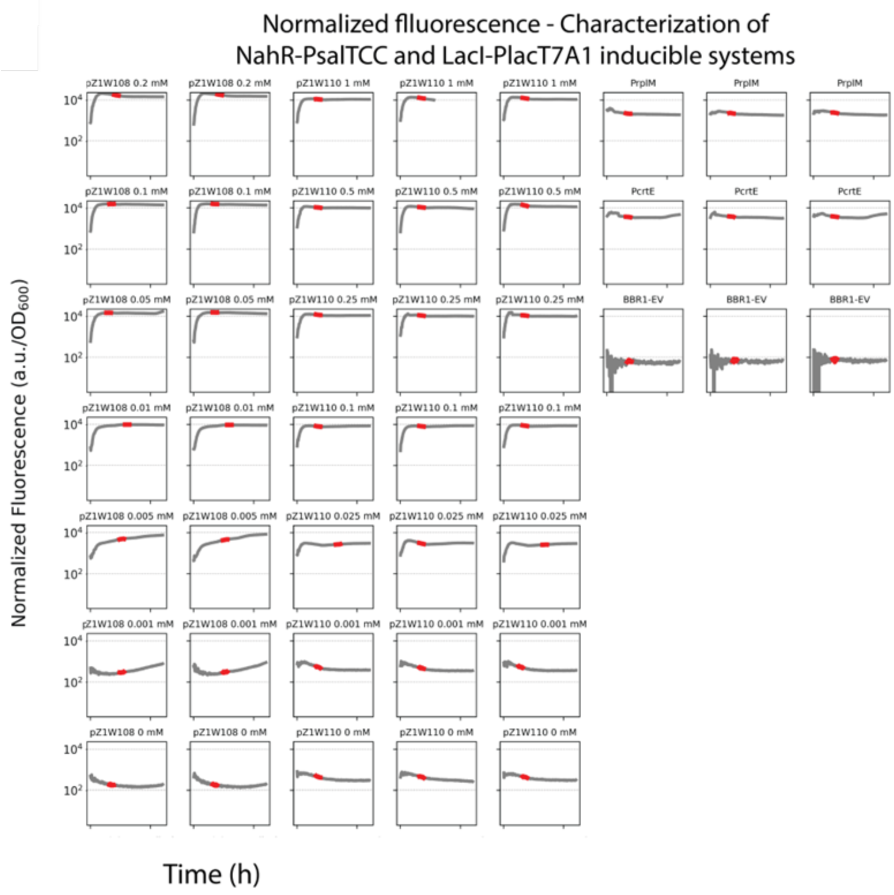


*Figure S5: Biomass-normalized fluorescence values of the assessment of inducible systems NahR-PsalTCC and LacI-PlacT7A_O3O4. Each plot represents a well in a 96-well plate reader. The highest moving average of 15 points during the exponential phase was used for the calculation of the average biomass-normalized fluorescence intensities and is highlighted in red.*

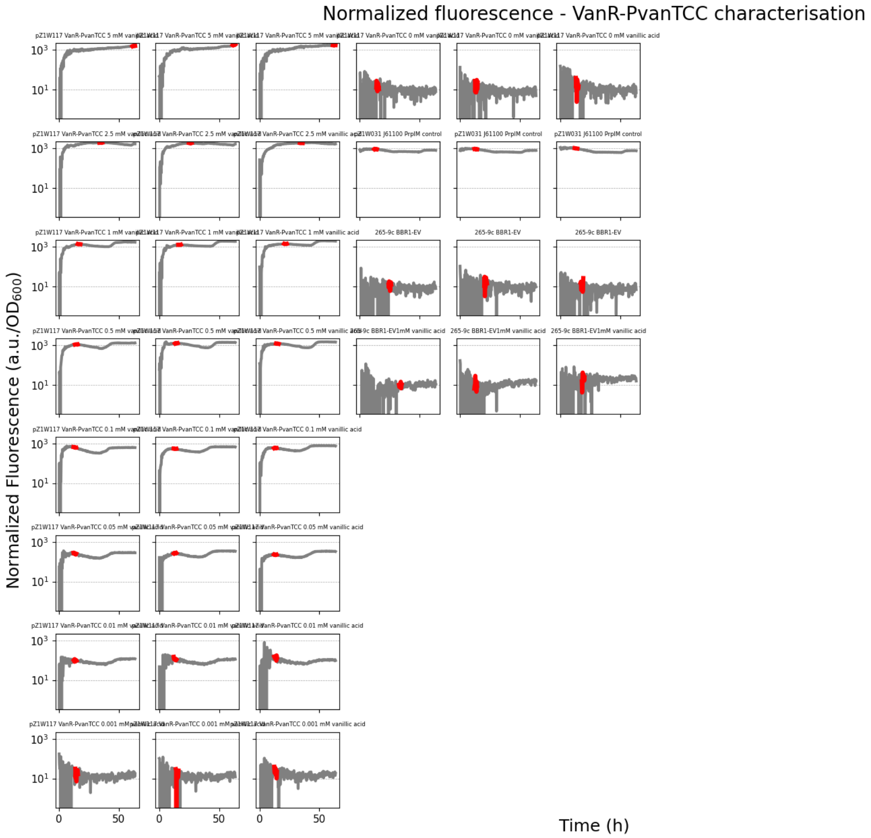


*Figure S6: Biomass-normalized fluorescence values of the assessment of inducible system VanR-PvanCC. Each plot represents a well in a 96-well plate reader. The highest moving average of 15 points during the exponential phase was used for the calculation of the average biomass-normalized fluorescence intensities and is highlighted in red.*

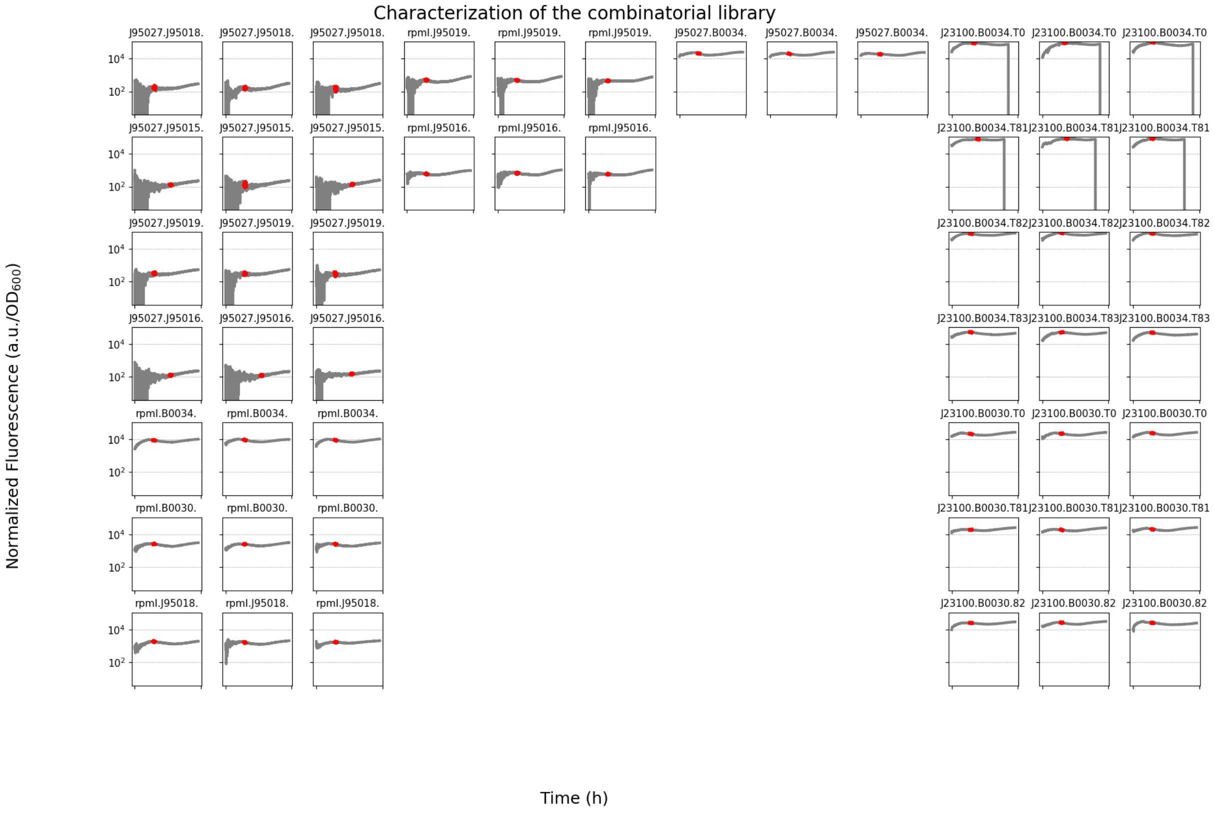


*Figure S7: Biomass-normalized fluorescence values of the combinatorial study. Each plot represents a well in a 96-well plate reader. The highest moving average of 15 points during the exponential phase was used for the calculation of the average biomass-normalized fluorescence intensities and is highlighted in red.*

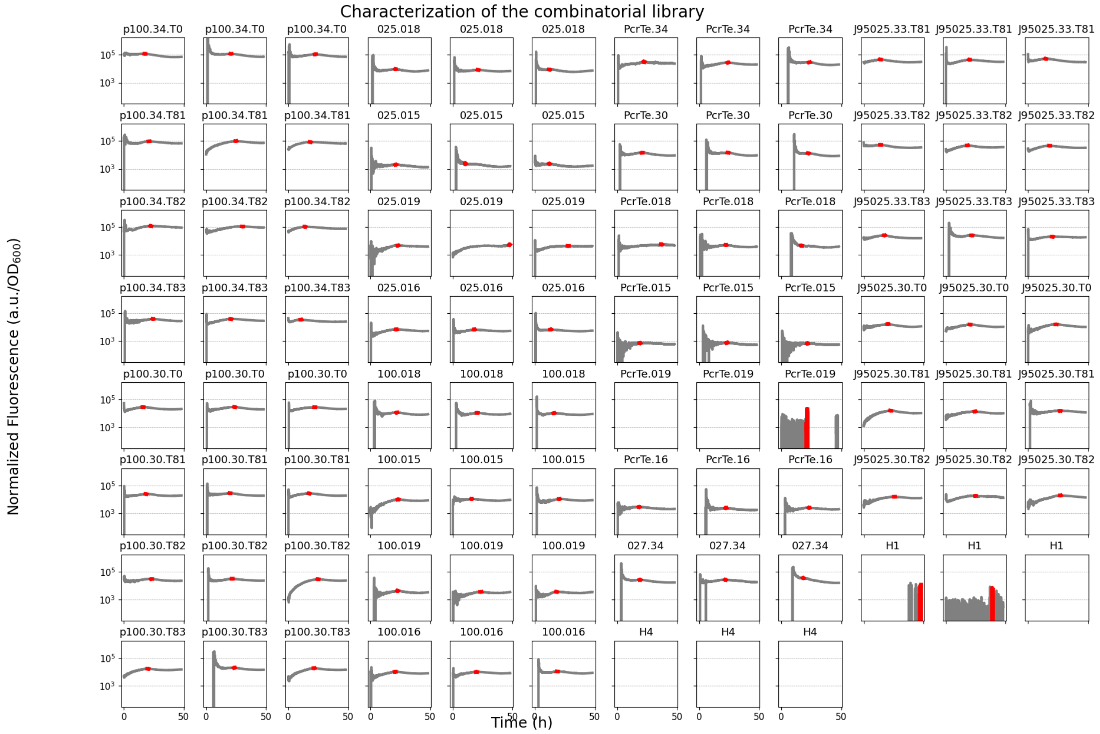


*Figure S14 Biomass-normalized fluorescence values of the combinatorial study. Each plot represents a well in a 96-well plate reader. The highest moving average of 15 points during the exponential phase was used for the calculation of the average biomass-normalized fluorescence intensities and is highlighted in red.*

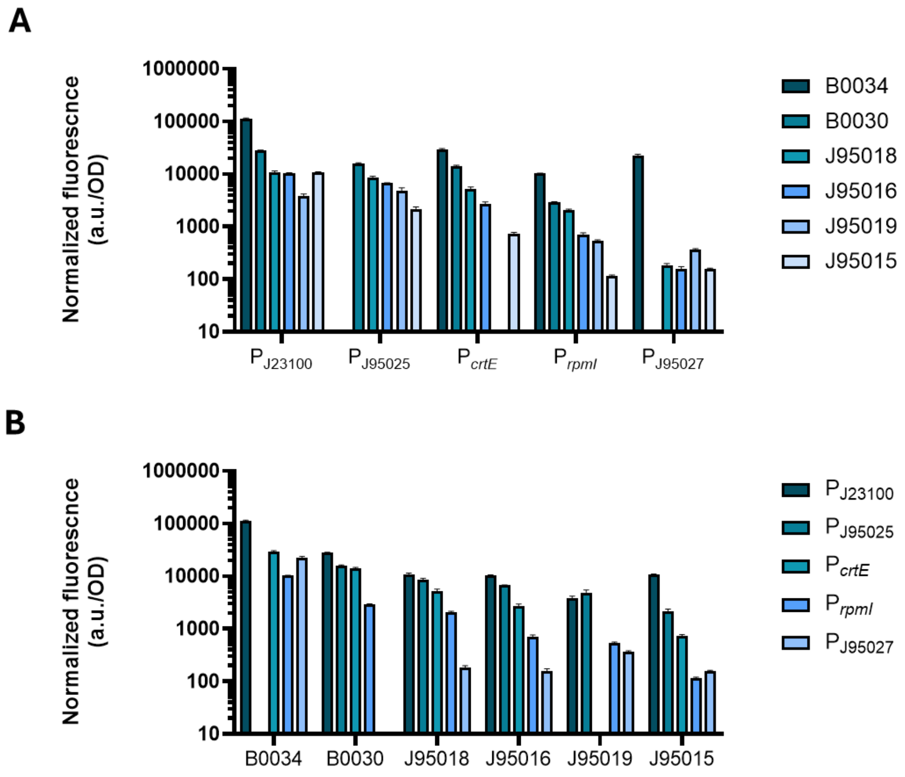


*Figure S15: Combinatorial promoter/RBS parts characterization. Average and standard deviation of the biomass-normalized fluorescence intensity are represented. A) The six tested RBSs were compared for each promoter. B) The five tested promoters are compared for each RBS.*

### Tables

*Table 1: Strains utilized in the study.*

| Strain | Source |
|---|---|
| R. sphaeroides 265-9c | (Orsi et al., 2019) |
| E. coli DH5α | New England Biolabs |
| E. coli ST18 | (Thoma & Schobert, 2009) |

*Table 2 : Plasmids utilized in the study.*

| Level 0 plasmids | | | | |
|---|---|---|---|---|
| ID | Level | Position | Part name | Source |
| pZ0W012 | 0 | 2 | J95015 | Huo (2011 |
| pZ0W013 | 0 | 2 | J95016 | Huo (2011) |
| pZ0W014 | 0 | 2 | J95017 | Huo (2011) |
| pZ0W015 | 0 | 2 | J95018 | Huo (2011) |
| pZ0W016 | 0 | 2 | J95021 | Huo (2011) |
| pZ0W017 | 0 | 2 | J95028 | Huo (2011) |
| pZ0W018 | 0 | 1 | $P_{J95027}$ | Huo (2011) |
| pZ0W019 | 0 | 1 | $P_{J23100}$ | Anderson (2006) |
| pZ0W020 | 0 | 1 | $P_{J23116}$ | Anderson (2006) |
| pZ0W024 | 0 | 1 | $P_{rpmI}$ | This study |
| pZ0W025 | 0 | 1 | $P_{crtE}$ | Beekwilder et al., 2014) |
| pZ0W026 | 0 | 4 | T0 | Lutz et al., (1997) |
| pZ0W027 | 0 | 2 | B0030 | Mahajan et al. (2003) |
| pZ0W029 | 0 | 1 | $P_{J23105}$ | Anderson (2006) |
| pZ0W031 | 0 | 2 | B0031 | Mahajan et al. (2003) |
| pZ0W032 | 0 | 2 | B0032 | Mahajan et al. (2003) |
| pZ0W033 | 0 | 2 | B0034 | Mahajan et al. (2003) |
| pZ0W034 | 0 | 2 | J61100 | Anderson (2007) |
| pZ0W035 | 0 | 1 | $P_{J23114}$ | Anderson (2006) |
| pZ0W036 | 0 | 2 | J95019 | Huo (2011) |
| pZ0W037 | 0 | 1 | $P_{J23104*}$ | Anderson (2006) |
| pZ0W042 | 0 | 1 | $P_{J95025}$ | Huo (2011) |
| pZ0W043 | 0 | 1 | $P_{J95023}$ | Huo (2011) |
| pZ0W081 | 0 | 4 | B1010 | Huang (2007) |
| pZ0W082 | 0 | 4 | B1015 | Huang (2007) |
| pZ0W083 | 0 | 4 | J95029 | Huo (2011) |
| pZP1321 | 0 | 1 | VanR-$P_{vanCC}$ | Shuster and Reich (2021), Meyer et al. (2019) |
| pZP1320 | 0 | 1 | NahR-$P_{sal}$ | Shuster and Reich (2021), Meyer et al. (2019) |

| | | | | |
|---|---|---|---|---|
| pZP284 | 0 | 1 | XylS-$P_{m}$ | Shuster and Reich (2021), Mermod et al. (1986) |
| pZP286 | 0 | 1 | LacI-$P_{lacT7A1}$ | Shuster and Reich (2021), Deuschle et al. (1986) |
| pZP434 | 0 | 1 | $P_{strong1k}$ | Behrendt et al. (2022) |
| pZP435 | 0 | 1 | $P_{strong10k}$ | Behrendt et al. (2022) |
| pZP436 | 0 | 1 | $P_{strong100k*}$ | Behrendt et al. (2022) |
| pZP124 | 0 | 3 | eGFP | Behrendt et al. (2022) |
| pZP001 | 0 entry (P) | Entry promoter | | Behrendt et al. (2022) |
| pZP002 | 0 entry (RBS) | Entry RBS | | Behrendt et al. (2022) |
| pZP003 | 0 entry (GOI) | Entry GOI | | Behrendt et al. (2022) |
| pZP004 | 0 entry (T) | Entry terminator | | Behrendt et al. (2022) |
| pZP1126 | 1 entry | Entry BBR1-KnR | | Behrendt et al. (2022) |
| pZP1125 | 1 entry | Entry RK2-KnR | | Behrendt et al. (2022) |
| pZP1395 | 1 entry | Entry RSF1010-KnR | | Behrendt et al. (2022) |

Level 1 plasmids (TUs)

| | ORI | Promoter | RBS | GOI | Terminator |
|---|---|---|---|---|---|
| pZ1W019 | pBBR1 | PrpmI | J95016 | EGFP | TO |
| pZ1W020 | pBBR1 | PrpmI | J95017 | EGFP | TO |
| pZ1W021 | pBBR1 | PrpmI | J95018 | EGFP | TO |
| pZ1W022 | pBBR1 | Pcrte | J95016 | EGFP | TO |
| pZ1W023 | pBBR1 | Pcrte | J95017 | EGFP | TO |
| pZ1W024 | pBBR1 | Pcrte | J95018 | EGFP | TO |
| pZ1W029 | pBBR1 | PrpmI | B0032 | EGFP | TO |
| pZ1W030 | pBBR1 | PrpmI | B0034 | EGFP | TO |
| pZ1W031 | pBBR1 | PrpmI | J61100 | EGFP | TO |
| pZ1W032 | pBBR1 | PrpmI | RBS_15 | EGFP | TO |
| pZ1W033 | pBBR1 | PrpmI | RBS_16 | EGFP | TO |
| pZ1W034 | pBBR1 | PrpmI | RBS_17 | EGFP | TO |
| pZ1W035 | pBBR1 | PrpmI | RBS_20 | EGFP | TO |
| pZ1W036 | pBBR1 | PrpmI | J95019 | EGFP | TO |
| pZ1W038 | pBBR1 | Pcrte | J95028 | EGFP | TO |
| pZ1W039 | pBBR1 | Pcrte | B0030 | EGFP | TO |
| pZ1W040 | pBBR1 | Pcrte | B0031 | EGFP | TO |
| pZ1W041 | pBBR1 | Pcrte | B0032 | EGFP | TO |
| pZ1W042 | pBBR1 | Pcrte | B0034 | EGFP | TO |
| pZ1W043 | pBBR1 | Pcrte | J61100 | EGFP | TO |
| pZ1W044 | pBBR1 | Pcrte | RBS_15 | EGFP | TO |
| pZ1W045 | pBBR1 | Pcrte | RBS_16 | EGFP | TO |
| pZ1W046 | pBBR1 | Pcrte | RBS_17 | EGFP | TO |
| pZ1W047 | pBBR1 | Pcrte | RBS_20 | EGFP | TO |
| pZ1W048 | pBBR1 | Pcrte | J95019 | EGFP | TO |
| pZ1W049 | pBBR1 | J95027 | J95017 | EGFP | TO |

| | | | | | |
|---|---|---|---|---|---|
| pZ1W050 | pBBR1 | pSppa | J95017 | EGFP | TO |
| pZ1W051 | pBBR1 | pStrong 1K | J95017 | EGFP | TO |
| pZ1W052 | pBBR1 | pStrong 10K | J95017 | EGFP | TO |
| pZ1W053 | pBBR1 | pStrong 100K* | J95017 | EGFP | TO |
| pZ1W054 | pBBR1 | J23100 | J95017 | EGFP | TO |
| pZ1W055 | pBBR1 | J23105 | J95017 | EGFP | TO |
| pZ1W056 | pBBR1 | J23116 | J95017 | EGFP | TO |
| pZ1W057 | pBBR1 | J23114 | J95017 | EGFP | TO |
| pZ1W058 | pBBR1 | J23104* | J95017 | EGFP | TO |
| pZ1W062 | pBBR1 | J95025 | J95017 | EGFP | TO |
| pZ1W063 | pBBR1 | J95023 | J95017 | EGFP | TO |
| pZ1W064 | pBBR1 | J95027 | J61100 | EGFP | TO |
| pZ1W065 | pBBR1 | pSppa | J61100 | EGFP | TO |
| pZ1W066 | pBBR1 | pStrong 1K | J61100 | EGFP | TO |
| pZ1W067 | pBBR1 | pStrong 10K | J61100 | EGFP | TO |
| pZ1W068 | pBBR1 | pStrong 100K* | J61100 | EGFP | TO |
| pZ1W069 | pBBR1 | J23100 | J61100 | EGFP | TO |
| pZ1W070 | pBBR1 | J23105 | J61100 | EGFP | TO |
| pZ1W071 | pBBR1 | J23116 | J61100 | EGFP | TO |
| pZ1W072 | pBBR1 | J23114 | J61100 | EGFP | TO |
| pZ1W073 | pBBR1 | J23104* | J61100 | EGFP | TO |
| pZ1W074 | pBBR1 | prsp_7571 | J61100 | EGFP | TO |
| pZ1W075 | pBBR1 | T334-25 | J61100 | EGFP | TO |
| pZ1W077 | pBBR1 | J95025 | J61100 | EGFP | TO |
| pZ1W078 | pBBR1 | J95023 | J61100 | EGFP | TO |
| pZ1W079 | RK2 | Rplm | J95017 | EGFP | TO |
| pZ1W081 | RSF1010 | Rplm | J95017 | EGFP | TO |
| pZ1W107 | pBBR1 | VanR-PvanCC (pZP1321) | J61100 | EGFP | TO |
| pZ1W108 | pBBR1 | NahR-Psal (pZP1320) | J61100 | EGFP | TO |
| pZ1W109 | pBBR1 | XylS-PM(pZP284) | J61100 | EGFP | TO |
| pZ1W110 | pBBR1 | LacI-Plact7Al (pZP286) | J61100 | EGFP | TO |
| pZ1WA001 | pBBR1 | PJ23100 | B0034 | EGFP | TO |
| pZ1WA002 | pBBR1 | PJ23100 | B0030 | EGFP | TO |
| pZ1WA003 | pBBR1 | PJ23100 | J95018 | EGFP | TO |
| pZ1WA004 | pBBR1 | PJ23100 | J95016 | EGFP | TO |
| pZ1WA005 | pBBR1 | PJ23100 | J95019 | EGFP | TO |
| pZ1WA006 | pBBR1 | PJ23100 | J95015 | EGFP | TO |
| pZ1WA007 | pBBR1 | PJ95025 | B0034 | EGFP | TO |
| pZ1WA008 | pBBR1 | PJ95025 | B0030 | EGFP | TO |
| pZ1WA009 | pBBR1 | PJ95025 | J95018 | EGFP | TO |
| pZ1WA010 | pBBR1 | PJ95025 | J95016 | EGFP | TO |
| pZ1WA011 | pBBR1 | PJ95025 | J95019 | EGFP | TO |
| pZ1WA012 | pBBR1 | PJ95025 | J95015 | EGFP | TO |

| | | | | | |
|---|---|---|---|---|---|
| pZ1WA013 | pBBR1 | Pcrte | B0034 | EGFP | TO |
| pZ1WA014 | pBBR1 | Pcrte | B0030 | EGFP | TO |
| pZ1WA015 | pBBR1 | Pcrte | J95018 | EGFP | TO |
| pZ1WA016 | pBBR1 | Pcrte | J95016 | EGFP | TO |
| pZ1WA017 | pBBR1 | Pcrte | J95019 | EGFP | TO |
| pZ1WA018 | pBBR1 | Pcrte | J95015 | EGFP | TO |
| pZ1WA019 | pBBR1 | Prplm | B0034 | EGFP | TO |
| pZ1WA020 | pBBR1 | Prplm | B0030 | EGFP | TO |
| pZ1WA021 | pBBR1 | Prplm | J95018 | EGFP | TO |
| pZ1WA022 | pBBR1 | Prplm | J95016 | EGFP | TO |
| pZ1WA023 | pBBR1 | Prplm | J95019 | EGFP | TO |
| pZ1WA024 | pBBR1 | Prplm | J95015 | EGFP | TO |
| pZ1WA025 | pBBR1 | PJ95027 | B0034 | EGFP | TO |
| pZ1WA026 | pBBR1 | PJ95027 | B0030 | EGFP | TO |
| pZ1WA027 | pBBR1 | PJ95027 | J95018 | EGFP | TO |
| pZ1WA028 | pBBR1 | PJ95027 | J95016 | EGFP | TO |
| pZ1WA029 | pBBR1 | PJ95027 | J95019 | EGFP | TO |
| pZ1WA030 | pBBR1 | PJ95027 | J95015 | EGFP | TO |
| pZ1WA031 | pBBR1 | PJ23100 | B0034 | EGFP | TO |
| pZ1WA032 | pBBR1 | PJ23100 | B0034 | EGFP | B1010 |
| pZ1WA033 | pBBR1 | PJ23100 | B0034 | EGFP | B1015 |
| pZ1WA034 | pBBR1 | PJ23100 | B0034 | EGFP | J95029 |
| pZ1WA035 | pBBR1 | PJ23100 | B0030 | EGFP | TO |
| pZ1WA036 | pBBR1 | PJ23100 | B0030 | EGFP | B1010 |
| pZ1WA037 | pBBR1 | PJ23100 | B0030 | EGFP | B1015 |
| pZ1WA038 | pBBR1 | PJ23100 | B0030 | EGFP | J95029 |
| pZ1WA039 | pBBR1 | PJ95025 | B0034 | EGFP | TO |
| pZ1WA040 | pBBR1 | PJ95025 | B0034 | EGFP | B1010 |
| pZ1WA041 | pBBR1 | PJ95025 | B0034 | EGFP | B1015 |
| pZ1WA042 | pBBR1 | PJ95025 | B0034 | EGFP | J95029 |
| pZ1WA043 | pBBR1 | PJ95025 | B0030 | EGFP | TO |
| pZ1WA044 | pBBR1 | PJ95025 | B0030 | EGFP | B1010 |
| pZ1WA045 | pBBR1 | PJ95025 | B0030 | EGFP | B1015 |
| pZ1WA046 | pBBR1 | PJ95025 | B0030 | EGFP | J95029 |

Broad-host expression vectors

| ID | Level | Positon | ORI | AbR | Dropout | Source |
|---|---|---|---|---|---|---|
| pSEVAb63-EV | SEVA vector | / | / | / | / | (Silva-Rocha et al., 2013) |
| pZP311 | 2 entry | 1 | p15A | SpecR | LacZ | Behrendt et al. (2022) |
| pZP334 | 1 entry | 4 | p15A | CamR | | Behrendt et al. (2022) |
| pZP1126 | 1 entry | 4 | pBBR1 | KnR | LacZ | Behrendt et al. (2022) |
| pZP1125 | 1 entry | 4 | RK2 | KnR | LacZ | Behrendt et al. (2022) |
| pZP1395 | 1 entry | 4 | RSF1010 | KnR | LacZ | Behrendt et al. (2022) |

| | | | | | | |
|---|---|---|---|---|---|---|
| pZ1W-E004 | 1 entry | 4 | RK2 | SpecR | LacZ | This study |
| pZ1W-E005 | 1 entry | 4 | RK2 | CamR | LacZ | This study |
| pZ1W-E006 | 1 entry | 4 | RK2 | GmR | LacZ | This study |
| pZ1W-E007 | 1 entry | 4 | RSF1010 | SpecR | LacZ | This study |
| pZ1W-E008 | 1 entry | 4 | RSF1010 | CamR | LacZ | This study |
| pZ1W-E009 | 1 entry | 4 | RSF1010 | GmR | LacZ | This study |
| pZ1W-E010 | 1 entry | 4 | pBBR1 | SpecR | LacZ | This study |
| pZ1W-E011 | 1 entry | 4 | pBBR1 | CamR | LacZ | This study |
| pZ1W-E012 | 1 entry | 4 | pBBR1 | GmR | LacZ | This study |
| pZ2W-E001 | 2 entry | 1 | RK2 | SpecR | mCherry | This study |
| pZ2W-E002 | 2 entry | 1 | RK2 | CamR | mCherry | This study |
| pZ2W-E003 | 2 entry | 1 | RK2 | GmR | mCherry | This study |
| pZ2W-E004 | 2 entry | 1 | RSF1010 | SpecR | mCherry | This study |
| pZ2W-E005 | 2 entry | 1 | RSF1010 | CamR | mCherry | This study |
| pZ2W-E006 | 2 entry | 1 | RSF1010 | GmR | mCherry | This study |
| pZ2W-E007 | 2 entry | 1 | pBBR1 | SpecR | mCherry | This study |
| pZ2W-E008 | 2 entry | 1 | pBBR1 | CamR | mCherry | This study |
| pZ2W-E009 | 2 entry | 1 | pBBR1 | GmR | mCherry | This study |

*Table S3: Sequences of assessed genetic elements.*

| Part name | Part Sequence |
|---|---|
| J95015 | CATCAACGGAGG |
| J95016 | CCTGGGGGAGGG |
| J95017 | TCAGTGGAGGGA |
| J95018 | GGAGGAGGGGGC |
| J95021 | GAGCAGAGGAGA |
| J95028 | GGAGGGGAGGCA |
| J95027 | AGCCCAAAAAATCCGCTTGCGCCCGGGGCCGTCTGCTCCTAGAAACCGCTTCATGTGGAATTGTGAG CGCTCACAATTCCACA |
| J23100 | TTGACGGCTAGCTCAGTCCTAGGTACAGTGCTAGC |
| J23116 | TTGACAGCTAGCTCAGTCCTAGGGACTATGCTAGC |
| RBS15 | TTATAAGGAGG |
| RBS16 | TATTTAAGGGGG |
| RBS17 | ATTTAAGGCGG |
| PrpmI | CATGGATGGGCGTCAAGGGTCAAGATGTCAGAAGGTTCATCTTGCTCACGTCACTGTGAGGACTTTCAC GTCGTCGTGACTTGAGGAAAGGAAGGACTTATGATTGAGCGTTCAAGAAAATAATGCGCTTGCAGGCCA CTTGGGCTTACGATAAGGACACCGCTTCCGCGCACGCGTCCGGGCCACAAGTGGCATGCCCGGCCG TGCAGAGTCTCTGCAGGATGCAGGACCAATCGTTTACGGCGAGGCAGACCCAT |
| PcrtE | GCTGCTGAACGCGATGGCGGCGCGGGGCGCGACGCGCGGGGCCGCATCCGTCTGCATCGGCGGG GGCGAGGCGACGGCCATCGCGCTGGAACGGCTGAGCTAATTCATTTGCGCGAATCCGCGTTTTTCGT GCACGATGGGGGAACCGGAAACGGCCACGCCTGTTGTGGTTGCGTCGACCTCTCTTCGGGCCATGC CCGTGACGCGATGTGGCAGGCGCATGGGGCGTTGCCGATCCGGTCGCATGACTGACGCAACGAAGG CACAT |
| TO | GACTCCTGTTGATAGATCCAGTAATGACCTCAGAACTCCATCTGGATTTGTTCAGAACGCTCGGTTGCCG CCGGGCGTTTTTTATTGGTGAGAAT |
| B0030 | AAAGAGGAGAAA |
| RBS20 | ATTTAAGGCTG |
| J23105 | TTTACGGCTAGCTCAGTCCTAGGTACTATGCTAGC |
| B0031 | TCTAGAGTCACACAGGAAACCTACTAG |
| B0032 | TCTAGAGTCACACAGGAAAGTACTAG |
| B0034 | TCTAGAGAAAGAGGAGAAATACTAG |
| J61100 | TCTAGAGAAAGAGGGGACAAACTAG |
| J23114 | TTTATGGCTAGCTCAGTCCTAGGTACAATGCTAGC |
| J95019 | TCGGAGGAGCCT |
| J23104* | TTGACAGCTAGCTCAGTCCTAGGTATTGTGCTAGC |
| J95025 | AAATTGTTACGGAGCCCAAAAAATCCGCTTGCGCCCGGGGCCGTCTGCTCCTAGAAACCGCTTCACC GAGACGTAGACCGGCAGCGCCGGACGGAGACGAGGGAGCGGATGACAGAAACGTCGGCCGCGAC AATTGAAGATGAGGCGGACGGGATCGCTGGTTGTCTG |
| J95023 | TCGTCTCTTCGTCATTTTTCCTCTTGCGGGTTTTTTTGCGGTTCCCTAGATAGCGCCTCACCGAAGCGGA ACGGCGACGGTGACGGGGTTGAGAGGCGGCGGTGCTGCCTTGAGGCTTTCGGAAATCTGGAAGATGA GGCGGACGGGATCGCTGGTT |
| B1010 | CCAGGCATCAAATAAAACGAAAGGCTCAGTCGAAAGACTGGGCCTTTCGTTTTATCTGTTGTTTGTCGGT GAACGCTCTC |
| B1015 | CCAGGCATCAAATAAAACGAAAGGCTCAGTCGAAAGACTGGGCCTTTCGTTTTATCTGTTGTTTGTCGGT GAACGCTCTCTACTAGAGTCACACTGGCTCACCTTCGGGTGGGCCTTTCTGCGTTTATA |

| | |
|---|---|
| J95029 | GATCCGGTGGATGACCTTTTGAATGACCTTTAATAGATTATATTACTAATTAATTGGGGACCCTAGAGGTC CCCTTTTTTATTTTAAAAATTTTTTCACAAAACGGTTTACAAGCATAAAGCTTGCTCAATCAATCACC |
| VanR-PvanCC | ATTGGATCCAATTGACAGCTAGCTCAGTCCTAGGTACCATTGGATCCAAT |
| NahR-Psal | GGGGCCTCGCTTGGGTTATTGCTGGTGCCCGGCCGGGCGCAATATTCATGTTGATGATTTATTATATATC GAGTGGTGTATTTATTTATATTGTTTGCTCCGTTACCGTTATTAAC |
| XylS-PM | TGCAAGAAGCGGATACAGGAGTGCAAAAAATGGCTATCTCTAGAATAGCCTACCCATTAGGCTTTATCAA CA |
| LacI-Plact7A1 | TTGACTTGTGAGCGGATAACAATGATACTTAGATTCAATTGTGAGCGGATAACAATT |
| EGFP | ATGGTGAGCAAGGGCGAGGAGCTGTTCACCGGGGTGGTGCCCATCCTGGTCGAGCTGGACGGCGAC GTAAACGGCCACAAGTTCAGCGTGCGCGGCGAGGGCGAGGGCGATGCCACCAACGGCAAGCTGAC CCTGAAGTTCATCTGCACCACCGGCAAGCTGCCCGTGCCCTGGCCCACCCTCGTGACCACCCTGAC CTACGGCGTGCAGTGCTTCAGCCGCTACCCCGACCACATGAAGCAGCACGACTTCTTCAAGTCCGCC ATGCCCGAAGGCTACGTCCAGGAGCGCACCATCAGCTTCAAGGACGACGGCACCTACAAGACCCGC GCCGAGGTGAAGTTCGAGGGCGACACCCTGGTGAACCGCATCGAGCTGAAGGGCATCGACTTCAAG GAGGACGGCAACATCCTGGGGCACAAGCTGGAGTACAACTTCAACAGCCACAACGTCTATATCACCG CCGACAAGCAGAAGAACGGCATCAAGGCCAACTTCAAGATCCGCCACAACGTGGAGGACGGCAGCG TGCAGCTCGCCGACCACTACCAGCAGAACACCCCCATCGGCGACGGCCCCGTGCTGCTGCCCGAC AACCACTACCTGAGCACCCAGTCCGTGCTGAGCAAAGACCCCAACGAGAAGCGCGATCACATGGTC CTGCTGGAGTTCGTGACCGCCGCCGGGATCACTCTCGGCATGGACGAGCTGTACAAGTAA |

*Table S4: Oligonucleotides used in this study*

| ID | Name | Sequence | Use |
|---|---|---|---|
| PR327 | pGGE-lvl0-seq_Fw | AGCGAGTCAGTGAGCGAG | primer |
| PR328 | pGGE-lvl0-seq_Rv | AGACGGTCACAGCTTGTCTG | primer |
| PR338 | 02_oligo_RBS_BBa_J95015_a | TAAGCAGAAGACATAAAACATCAACGGAGGAATGATGTCTTCTAAGCA | oligo |
| PR339 | 02_oligo_RBS_BBa_J95015_b | TGCTTAGAAGACATCATTCCTCCGTTGATGTTTTATGTCTTCTGCTTA | oligo |
| PR340 | 02_oligo_RBS_BBa_J95016_a | TAAGCAGAAGACATAAAACCTGGGGGAGGGAATGATGTCTTCTAAGCA | oligo |
| PR341 | 02_oligo_RBS_BBa_J95016_b | TGCTTAGAAGACATCATTCCCTCCCCCAGGTTTTATGTCTTCTGCTTA | oligo |
| PR342 | 02_oligo_RBS_BBa_J95017_a | TAAGCAGAAGACATAAAATCAGTGGAGGGAAATGATGTCTTCTAAGCA | oligo |
| PR343 | 02_oligo_RBS_BBa_J95017_b | TGCTTAGAAGACATCATTTCCCTCCACTGATTTTATGTCTTCTGCTTA | oligo |
| PR344 | 02_oligo_RBS_BBa_J95018_a | TAAGCAGAAGACATAAAAGGAGGAGGGGGCAATGATGTCTTCTAAGCA | oligo |
| PR345 | 02_oligo_RBS_BBa_J95018_b | TGCTTAGAAGACATCATTGCCCCCTCCTCCTTTTATGTCTTCTGCTTA | oligo |
| PR346 | 02_oligo_RBS_BBa_J95019_a | TAAGCAGAAGACATAAAATCGGAGGAGCCTAATGATGTCTTCTAAGCA | oligo |
| PR347 | 02_oligo_RBS_BBa_J95019_b | TGCTTAGAAGACATCATTAGGCTCCTCCGATTTTATGTCTTCTGCTTA | oligo |
| PR348 | 02_oligo_RBS_BBa_J95021_a | TAAGCAGAAGACATAAAAGAGCAGAGGAGAAATGATGTCTTCTAAGCA | oligo |
| PR349 | 02_oligo_RBS_BBa_J95021_b | TGCTTAGAAGACATCATTTCTCCTCTGCTCTTTTATGTCTTCTGCTTA | oligo |
| PR350 | 02_oligo_RBS_BBa_J95028_a | TAAGCAGAAGACATAAAAGGAGGGGAGGCAAATGATGTCTTCTAAGCA | oligo |
| PR351 | 02_oligo_RBS_BBa_J95028_b | TGCTTAGAAGACATCATTTGCCTCCCCTCCTTTTATGTCTTCTGCTTA | oligo |
| PR352 | 01_oligo_BBaJ95027_a_a | ATATAGAAGACATAGTGAGCCCAAAAAATCCGCTTGCGCCCGGGGCCGTCTGCTATGTCTTCTAAGCA | oligo |
| PR353 | 01_oligo_BBaJ95027_a_b | TGCTTAGAAGACATAGCAGACGGCCCCGGGCGCAAGCGGATTTTTTGGGCTCACTATGTCTTCTATAT | oligo |
| PR354 | 01_oligo_BBaJ95027_b_a | TAAGCAGAAGACATTGCTCCTAGAAACCGCTTCATGTGGAATTGTGAGCGCTCACAATTCCACAAAAAATGTCTTCAATCTCT | oligo |
| PR355 | 01_oligo_BBaJ95027_b_b | AGAGATTGAAGACATTTTTTGTGGAATTGTGAGCGCTCACAATTCCACATGAAGCGGTTTCTAGGAGCAATGTCTTCTGCTTA | oligo |
| PR356 | pZ1W_check_Fw | AGGTTGGGCTTCGGAATCG | primer |
| PR357 | pZ1W_check_Rv | GGTCACACTGCTTCCGGTAG | primer |
| PR372 | 01_pcr_Bba_J95023_Fw | ATATAGAAGACATAGTGTCGTCTCTTCGTCATTTTTCCTC | primer |
| PR373 | 01_pcr_Bba_J95023_Rv | GAAGACATTTTTAACCAGCGATCCCGTCC | primer |
| PR396 | RK2_lvl1_fw | CTATCAGGTCAAGTCTGCCCG | primer |
| PR397 | RK2_lvl1_rv | TGGACGATGGCCCACTCC | primer |
| PR456 | CamR_3'_SapI | GACTCAGCTCTTCATCCATTTAAATGGCGCGCCTTACG | primer |
| PR457 | CamR_5'SapI | GACTCAGCTCTTCATCGTGATCGGCACGTAAGAGGTTC | primer |
| PR458 | SpecR_3'_sapI_a_Fw | GACTCAGCTCTTCATCCCAGCTCTCTAACGCTTGAGTTAAG | primer |
| PR459 | SpecR_5' SapI_Rv | GACTCAGCTCTTCATCGCACCTGAAGTCAGCCCCATAC | primer |
| PR460 | GmR_5'_SapI | GACTCAGCTCTTCATCGGGGTCCCCAATAATTACGATTTAAATTTG | primer |
| PR461 | GmR_3'SapI | GACTCAGCTCTTCATCCCGGACCGTTGTCCAATTTACC | primer |
| PR462 | BB_Sap_Fw | GACTCAGCTCTTCACGACTGTCTCTTGATCAGATCTTGATCC | primer |
| PR463 | BB_sap_Rv | GACTCAGCTCTTCAGGAATGACCGACCAAGCGACG | primer |
| PR464 | Insert_lvl2pos1_SapI_Fw | GACTCAGCTCTTCATCCGGTCTCAAGTGAATGCCAAGTCTTCCTAGCTATCTACATATATATATATGTTGACAGGG | primer |

| | | | |
|---|---|---|---|
| PR465 | Insert_lvl2pos1_SapI_Rv | GACTCAGCTCTTCATCGGGTCTCACCTGAATCCTAAGTCTTCAAGATGAACAAAACTAAAGCGCCA | primer |
| PR466 | BB_1125_Sap_Fw | GACTCAGCTCTTCAGGACAGACTTGACCTGATAGTTTGGCTG | primer |
| PR467 | BB_1125_Sap_Rv | GACTCAGCTCTTCACGACGGGAGTGGGCCATCGTC | primer |
| PR468 | BB_1126_Sap_Fw | GACTCAGCTCTTCACGAGGCGAAGAACTCCAGCATGAG | primer |
| PR469 | BB_1126_Sap_Rv | GACTCAGCTCTTCAGGAAGCCGCTTATGTCTATTGCTGG | primer |
| PR470 | BB_1395_Sap_Rv | GACTCAGCTCTTCAGGATCCGGGTTTTTTTAAGGCAGT | primer |
| PR475 | B1006_a | CTACTAGGAAGACATCGAAAAAAAAAAAACCCCGCCCCTGACAGGGCGGGGTTTTTTTTAACCATGTCTTCTCTAGAGTC | oligo |
| PR476 | B1006_b | GACTCTAGAGAAGACATGGTTAAAAAAAACCCCGCCCTGTCAGGGGCGGGGTTTTTTTTTTTTCGATGTCTTCCTAGTAG | oligo |
| PR477 | B1002_a | CTACTAGGAAGACATCGAACGCAAAAAACCCCGCTTCGGCGGGGTTTTTTCGCAACCATGTCTTCTCTAGAGTC | oligo |
| PR478 | B1002_b | GACTCTAGAGAAGACATGGTTGCGAAAAAACCCCGCCGAAGCGGGGTTTTTTGCGTTCGATGTCTTCCTAGTAG | oligo |
| PR479 | B1003_a | CTACTAGGAAGACATCGAACGCCAAAAACCCCGCTTCGGCGGGGTTTTTCCGCAACCATGTCTTCTCTAGAGTC | oligo |
| PR480 | B1003_b | GACTCTAGAGAAGACATGGTTGCGGAAAAACCCCGCCGAAGCGGGGTTTTTGGCGTTCGATGTCTTCCTAGTAG | oligo |
