## Supporting Information 2 for "Rhodo-Box: a Synthetic Biology Toolbox to Facilitate Metabolic Engineering of *Rhodobacter sphaeroides*"

### Supplementary material and method :

#### Automated Golden Gate assembly

The automated Golden Gate assembly protocol was developed using the Opentrons Python API v2.0. An Excel file was used to indicate the desired assemblies to perform. This protocol required a P20 GEN2 pipette connected to the left arm of the OT2. Two modules were used in this protocol: a thermocycler GEN1 and a temperature module GEN2 with an Opentrons 24 aluminium block with NEST 1.5 mL Snapcap. Opentrons 96 20 µL filter tip racks were used along with two NEST 100 µL full skirt 96 well plates, one source plate loaded with plasmids and one empty destination plate loaded in the thermocycler.

The automated protocol entailed mixing 1 µL of different plasmids in the destination plate according to the excel file indications. The mixtures were diluted with Milli-Q water to a final volume of 7.5 µL. A master mix of the Golden Gate reaction described in the Material and Method section was stored on the temperature module at 10 º C. 2.5 µL of this master mix was finally added to each specific plasmid mixture in the thermocycler. 30 cycles of 5 min 37 ºC and 5 min 16 ºC were performed in the thermocycler, followed by a longer digestion step at 37 ºC for 10 min, before holding the temperature at 10 ºC until manual termination of the protocol.

#### Automated chemical transformation

The automated chemical transformation protocol was designed using the Opentrons protocol designer webpage. An HEPA module is required to work in sterile conditions for this protocol. A P20 GEN2 8 channels pipette was used on the left arm of the OT2 and a P300 GEN2 8 channels pipette loaded on the right arm of the OT2. The thermocycler module GEN2 was present from the previous protocol. The aluminium block of the temperature module GEN2 was switched to an Opentrons 96-well aluminium block with generic 200 µL PCR strips. Opentrons 96 20 and 200 µL filter tip racks were used along with NEST 12 15 mL well reservoirs. Well A1 was loaded with LB. 50 µL of chemically competent *E. coli* DH5a or ST18 were loaded on the temperature module, either in PCR strips or in a sterile NEST 96 100 µL well plates with full skirt.

The automated protocol entailed transferring the Golden Gate reaction from the thermocycler to the chemically competent *E. coli* cells cooled at 4 ºC on the temperature module. Following 20 min at 4 ºC, the temperature module was switched to 42 ºC for 45 s, before cooling back to 4 ºC for 20 min. 100 µL of LB was transferred to the cells and incubated for 1 h at 37 ºC.

The template provided is designed for 24 chemical transformations.

#### Automated plating

The automated chemical transformation protocol was designed using the Opentrons protocol designer webpage. This protocol required a HEPA module is required to work in sterile conditions. A P20 GEN2 8 channels pipette was loaded on the left arm of the OT2. An Opentrons 96 filter tip rack 20 µL labware was used, along with an SBS plate with 50 mL of LB agar, supplemented with the appropriate antibiotic for plasmid selection. The automated protocol entailed plating 5 µL of transformed cells onto the agar plate. Replicates were performed if the agar plate was not fully plated. Plating multiple SBS plates and varying dilutions was also performed by customizing this template protocol in the protocol designer page.

The template provided is designed for the plating of 24 transformed cells, in 4 replicates.

#### Supplementary Figure SI2

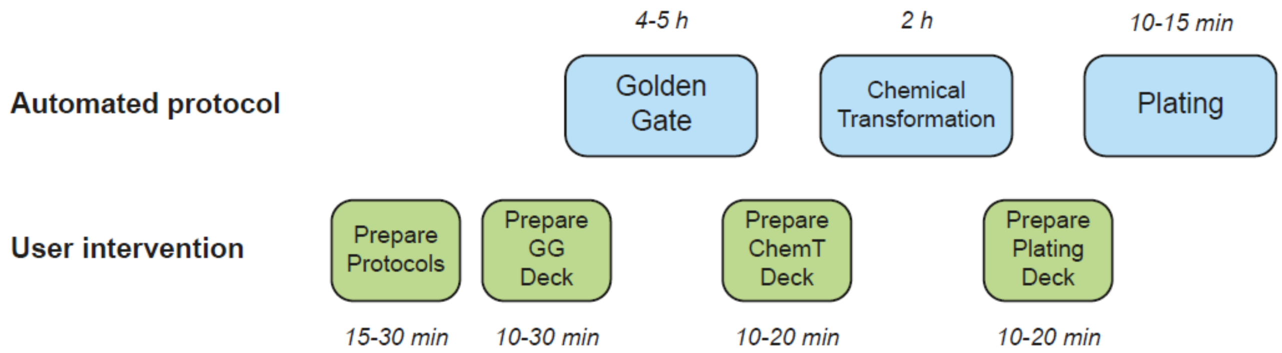


*Figure S21. General overview of the semi-automated cloning workflow. The automated physical protocols are represented in the blue frames, and the user interventions are indicated in the green frames. The approximate time for each step is indicated.*

#### Supplementary Table SI2

*Table S21. Overview of the digital and physical preparation steps for each individual protocol of the workflow.*

| | Golden Gate | Chemical transformation | Plating |
|---|---|---|---|
| **Digital protocols preparation** | Template excel :<br>• Fill source plate with parts to assemble<br>• Indicate assemblies in assembly plate<br>• Copy final parameters<br><br>Template python protocol :<br>• Paste parameters<br><br>Opentrons OT2 :<br>• Load the updated protocol | Opentrons protocol designer webpage :<br>• Import template protocol<br>• Edit protocol<br>• For each transfer step, add source and destination columns according to the experiment<br>• Rename, save, and export<br><br>Opentrons OT2 :<br>• Load the updated protocol | Opentrons protocol designer webpage :<br>• Import template protocol<br>• Edit protocol<br>• Add source and destination columns according to the experiment<br>• Rename, save, and export<br><br>Opentrons OT2 :<br>• Load the updated protocol |
| **Physical OT2 preparation** | • Load left P20 single (2-3 min)<br>• Load thermocycler (3 min)<br>• Load temperature module (2 min)<br>• Prepare source plate according to the excel (variable – 10 min)<br>• Load labware (5-10 min) :<br> • Pipette tips<br> • Reservoir $H_2O$<br> • Source plate<br> • Destination plate<br> • GG master mix | • Load P20 (left) and P300 (right) 8 channels (5 min)<br>• Load temperature module (2 min)<br>• Prepare competent cells on ice (3 min)<br>• Sterilize the OT2 and turn on the HEPA module (3 min)<br>• Load labware (5 min) :<br>• Pipette tips<br>• Reservoir with LB | • Load SBS agar plate<br>• Load pipette tips<br>(left P20 8 channel and temperature block with transformed cells already on deck) |

<u>Automation Supplementary Protocols :</u>

### Digital protocol preparation

#### Automated Golden Gate – *(10-20 min)*

1. Open “Template GG” excel file.
2. Fill one or the two source plates with the name of the parts to assemble. Do not use the symbol “_” in the names.

**Source plate 1**

| | 1 | 2 | 3 | 4 | 5 | 6 | 7 | 8 | 9 | 10 | 11 | 12 |
|---|---|---|---|---|---|---|---|---|---|---|---|---|
| A | ID0057 | ID448 | | | | | | | | | | |
| B | ID0058 | ID449 | | | | | | | | | | |
| C | ID0059 | ID450 | | | | | | | | | | |
| D | ID0060 | ID5001 | | | | | | | | | | |
| E | ID0061 | ID5002 | | | | | | | | | | |
| F | ID0062 | ID5003 | | | | | | | | | | |
| G | ID0063 | ID5004 | | | | | | | | | | |
| H | ID0064 | ID5005 | | | | | | | | | | |

3. A cell can be found with characters to copy (ctrl + v):
   *&"_"&*
   Do not copy the cell. Open the cell and copy the text.
4. The assembly plate should be filled in the following order: A1, B1, C1,… ,A2,B2, C2,… as shown below.

**Assembly plate**

| | 1 | 2 | 3 | 4 | 5 | 6 | 7 | 8 | 9 | 10 | 11 | 12 |
|---|---|---|---|---|---|---|---|---|---|---|---|---|
| A | 1 | 9 | | | | | | | | | | |
| B | 2 | 10 | | | | | | | | | | |
| C | 3 | 11 | | | | | | | | | | |
| D | 4 | … | | | | | | | | | | |
| E | 5 | | | | | | | | | | | |
| F | 6 | | | | | | | | | | | |
| G | 7 | | | | | | | | | | | |
| H | 8 | | | | | | | | | | | |

5. To indicate a specific assembly in a cell of the assembly plate, proceed as following :
   a. Start with “=” in the cell
   b. In the source plate, select and click on the part needed in this assembly. The excel coordinates will appear in the destination cell.
   c. Paste the characters previously copied (ctrl + v). This is a delimiter between each part of the assembly.
   d. Repeat operation b and c until all the parts have been indicated.
   e. Finish the cell assembly with coordinates, not the delimiter. Press enter.

| | 1 |
|---|---|
| A | =B3&"_"&B7&"_"&C5&"_"&C8 |
| B | |
| C | |
| D | |
| E | |
| F | |
| G | |
| H | |

| | 1 |
|---|---|
| A | ID0057_ID0061_ID450_ID5003 |
| B | |
| C | |
| D | |
| E | |
| F | |
| G | |
| H | |

6. Each assembly requires at least two parts.
7. Do not drag cells in the source or assembly plate. This would results in altered coordinates and references for the final parameters.

8. A column with the final parameters concatenated can be found in column AD: “Plate,Source_well,Destination_well,h2o”.
   Copy the column down to the last coordinates indicated.

| AD |
|---|
| Plate,Source_well,Destination_well,h2o |
| P1,A1,A1,3,5 |
| P1,E1,A1,5,5 |
| P1,C2,A1,4,5 |
| P1,F2,A1,4,5 |
| P1,F1,B1,4,5 |
| P1,E2,B1,3,5 |
| P1,A1,C1, |
| P1,F2,C1, |
| P1,G1,C1, |
| P1,G1,D1, |
| P1,B2,D1, |
| P1,H2,D1, |
| P1,F1,E1, |
| P1,C2,E1, |
| P1,G2,E1, |
| P1,B1,F1, |
| P1,E1,F1, |
| P1,G1,F1, |
| P1,C2,F1, |
| Not Found,,G1, |
| Not Found,,G1, |
| Not Found,,G1, |
| Not Found,,G1, |

9. Open the “Template Golden Gate” python file.
10. Paste the final parameters in the csv section at the top of the code as indicated.
11. Save and rename this specific protocol.
12. Load this specific protocol in the OT2 software.

#### **Automated chemical transformation** – *(5 min)*

1. Import the “Template Chemical Transformation” protocol in the opentrons designer web page.
2. Edit the protocol.
3. In the starting deck, add the adequate amount of assemblies present in the thermocycler, and a corresponding amount of competent cells on the thermoblock.

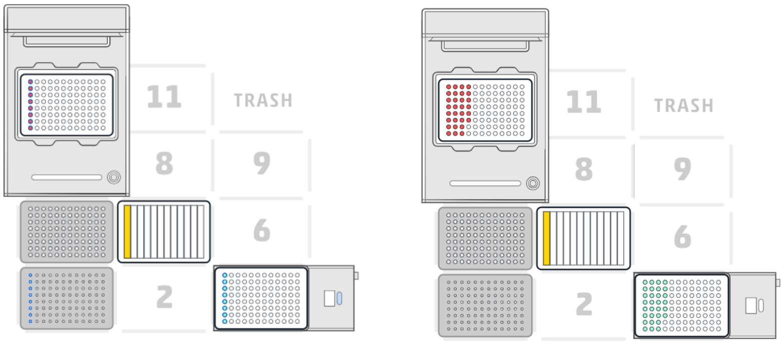


4. In the protocol steps, edit the two transfer steps (step 5 and step 15) to fit the amount of transformations to perform.
    a. Plasmids to competent cells

    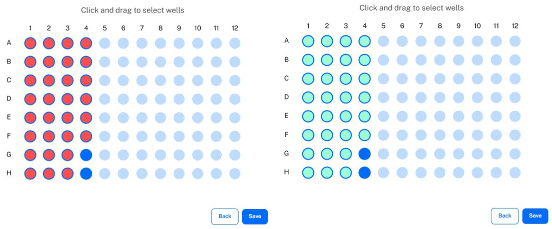
    

    *You can ignore the message about aspiring empty wells.*

    b. LB to competent cells
    The LB stays same well. Only indicate the changes for the destination in this transfer step.
5. Press Done at the top right of the edition page.
6. Rename the protocol.
7. Export the protocol.
8. Load the protocol in the OT2 software.

#### **Automated plating** – *(5 min)*

1. Import the plating template protocol in the opentrons designer web page.
2. Edit the protocol.
3. In the starting deck, add the adequate amount of transformed cells present on the temperature module.

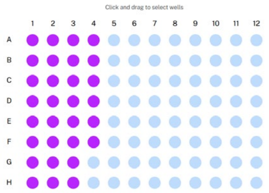


4. In the protocol steps, edit the only transfer step to fit the amount of plating to perform.

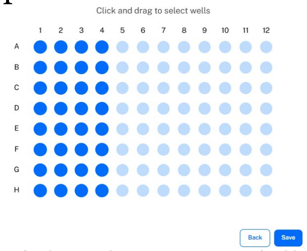


5. If the plate is not fully covered, plating replicates can be performed. Duplicate the transfer step multiple times until having as many transfer steps as columns. You will receive a tip error message that you can ignore for now.
    a. For the **first** transfer, select in the source plate the **first** column. In the destination plate, select the amount of replicates desired for this column. Change the tip handling to **"once at the start of the step".**

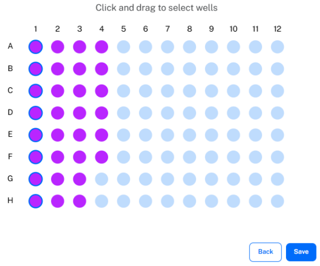


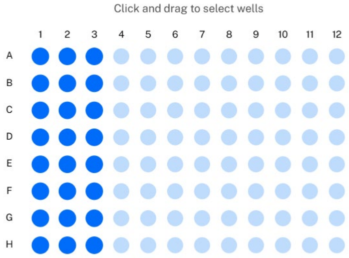


    b. For the **second** transfer, select in the source plate the **second** column. In the destination plate, select the amount of replicates desired for this column. Change the tip handling to **"once at the start of the step"**.

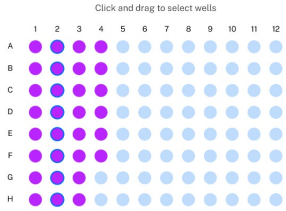


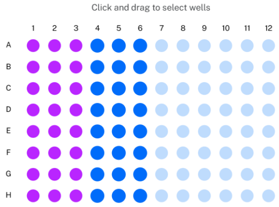


    c. Continue until the last transfer. Changing the tip handling should overcome the error message received at the start.

6. Press Done at the top right of the edition page.
7. Rename the protocol.

8. Export the protocol.
9. Load the protocol in the OT2 software.

### Physical protocol preparation

#### Automated Golden Gate – *(10-30 min)*

1. Prepare the Golden Gate Master-Mix with the adequate restriction enzyme.
2. Load the source plate with the same parts and layout as specified in the Excel assembly plate. 40 µL of a 50-100 ng/µL plasmid concentration should be used.
3. Load the pipette P20 single channel on the left arm of the OT2
4. Load the modules on the OT2:
   a. Thermocycler in position 7.
   b. Temperature module in position 6, with a 24 well aluminum block NEST 1.5 mL Snapcap adapter.
5. Load the labware and reagents on the deck:
   a. 20 µl tip boxes in positions 1, 4 and 9.
   b. 12 well reservoir in position 3, 10 mL of MiliQ water in well A1.
   c. On the temperature module, load the Golden Gate Master-Mix in position A1.
   d. Source plate 1 in position 2; Source plate 2 in position 5.
6. Start the protocol in the OT2 software.

#### Automated chemical transformation – *(10-20 min)*

1. Load pipettes P20 8 channels on the left arm and P300 8 channels on the right arm of the OT2.
2. The thermocycler and temperature module should be loaded from last protocol. If not, load them in position 7 and 6 respectively. Switch the temperature module adapter for a 96 well aluminum block.
3. Turn on the HEPA module and sterilize the OT2. Work sterile from now on.
4. Start the protocol when the modules are loaded, not necessarily the labware (the temperature module will cool down to 4 °C, taking some time for you to load the labware and the chemically competent cells. You'll need to resume manually once the deck is fully loaded).
5. Get chemically competent *E. coli* cells on ice next to the OT2.
6. Load the labware :
   a. Sterile P20 tips in position 1 and sterile P200 tips in position 4.
   b. Sterile 12 wells reservoir in position 5, 10 mL of LB in well A1.
   c. The source plate should be loaded in the thermocycler.
   d. Sterile 96 well plate loaded on the temperature module if competent cells not in PCR stripes.
7. Once the temperature block goes below 10 °C, load the chemically competent cells (from full 8 PCR strips or load a sterile 96 well plate).
8. Once the temperature module reaches 4 °C, a message will ask to resume the protocol when all the competent cells are loaded.

#### Automated plating – *(10-20 min)*

1. Prepare the SBS agar plate with the desired medium and antibiotic resistance.
   a. 50 mL of agar has to be used for efficient plating.
   b. The drying surface should be absolutely flat. Avoid using a flow cabinet with potentially bent surface. Use a spirit level if needed.

2. The pipette P20 8 channels should be loaded on the left arm of the OT2 from previous protocol.
3. The temperature module should be loaded with the transformed cells in position 6 from previous protocol.
4. The HEPA module should be ON from previous protocol. Work sterile in the OT2.
5. Load new sterile P20 tips on position 1.
6. Load the dried LB agar plate in position 2.
7. Start the protocol in the OT2 software.
